# Body size, wing area, and wing loading follow a pattern of trait conservatism in *Drosophila*

**DOI:** 10.64898/2026.02.12.705663

**Authors:** Jonathan A. Rader, Patrick W. Kelly, Gabriel A. Jimenez, Daniel R. Matute

## Abstract

Wing loading is a metric of flight performance that captures the relationship between body mass and wing area and reflects how much weight each unit of wing surface supports during flight. Comparative studies have documented substantial differences in wing loading among individuals and across species. However, no study has evaluated the extent of this variation when species are reared under identical, controlled conditions. Here, we address that gap by measuring wing loading in 30 species of drosophilids raised in a common lab environment. We applied comparative phylogenetic methods to assess the extent to which the evolution of body mass, wing area, and wing loading is structured by shared ancestry. We find that wing area and body mass exhibit moderate phylogenetic signal, but wing loading does not. In addition, all three traits are best explained by a model of evolution in which most trait divergence occurs during speciation events. More conservative analyses provide no support for adaptive peaks in wing loading within drosophilids. Together, our results indicate that the evolutionary dynamics of wing loading in *Drosophila* differ from those described in birds and bats, and raise the question of whether similar patterns characterize other insect lineages.

## INTRODUCTION

The performance of biomechanical systems determines how animals interact with their environment. The contribution of individual traits to the overall function of biomechanical systems mediates how natural selection acts on morphological variation associated with locomotion, feeding, and reproduction, providing the link between traits and the fitness that they confer [1]. This connection makes the dissection of the underpinnings, processes, and functional constraints that lead to biomechanical performance key for understanding how evolutionary processes shape form, function, and diversification across lineages.

Flight is the dominant mode of locomotion for numerous animal taxa, and the unique suite of traits that are associated with animal flight influences many other aspects of these animals’ biology and often exhibits close associations with fitness. For example, wing shape is classically thought to correlate with habitat and/or flight behavior and numerous other aspects of biology across most flying taxa [e.g., 2–6]. Relatively long, pointed wings are associated with longer migratory and dispersal distances in birds [3, 6, 7]. Similarly, both the size and shape of migratory species and populations differ from their sedentary counterparts in bats [8, 9] and a diverse array of insect taxa [10–12]. In tropical butterfly communities, species are stratified vertically through the forest canopy and wing shape differs among these microhabitats [13]. Alterations in wing shape (or reductions in wing area) either from feather molt in birds or ablative wing damage in insects typically result in reduced flight performance, manifesting in lower takeoff velocities, reduced maneuverability, and lower flight speeds [14–16], potentially reducing foraging effectiveness and survival [17, 18] [but see 19]. Furthermore, wing size is a direct predictor of fitness in *Trichogramma* parasitoid wasps [20].

Wing loading, one of the key determinants of flight performance, is the relationship between the body mass and the wing area of an individual and measures the amount of weight each unit of wing area supports during flight [21, 22]. All else being equal, fliers with lower wing loading may be able to produce a greater amount of aerodynamic force relative to their body mass, thus permitting greater accelerations than are possible when wing loading is greater [22, 23]. This may be important in escape flight, wherein the flier is using maximal performance to escape predation, or conversely, during hunting bouts, during which a predator must overcome an escape attempt. Larger wings, which have much greater virtual mass arising from the mass of air that they displace during acceleration [22], are useful for inertial reorientation during flight, increasing the maneuverability of the flier [24, 25], but this can be a double-edged sword. Increased wing mass also results in increased power requirement and energetic costs of flapping, potentially reducing the overall energetic efficiency of flight [22, 26, 27]. Low wing loading allows for maneuverability, facilitates the generation of lift, and is common in gliding and small birds. Conversely, high wing loading enables higher flight speeds and improved stability, but requires a greater energetic investment and is often observed in migratory and fast-flying species. Because wing loading is so intimately tied to flight performance, the traits that integrate wing morphology and flight performance with fitness and ecology may be especially susceptible to selection.

Comparative work has revealed extensive differences in wing loading among individuals and species. In butterflies and vertebrates, wing loading and flight speed show a positive correlation [28]. In wind-dispersed seeds, wing loading predicts the rate of descent after detaching from the tree, and as a result, the mean dispersal distance of the seeds can vary by almost an order of magnitude between species [29]. More generally, wing loading has important consequences for geographic range size [30], population structure [31], and species community composition [32], and it is correlated with species vulnerability to habitat fragmentation [33]. While there is evidence of interspecific differences among insect species [34], to date, no systematic effort has catalogued the scale of wing loading in a phylogenetically-informed perspective while controlling for developmental conditions and age.

The genus *Drosophila* represents a powerful model to study phenotypic evolution for at least two reasons. First, the genus is globally distributed and contains over 2,000 described species [35]. Second, *Drosophila* species have diversified to occupy a variety of ecological niches and every continent, with the exception of Antarctica. During this radiation, drosophilids diverged in their morphology, physiology, and behavior [35]. For example, thorax length, a proxy of body size, ranges from 0.66 mm in *Scaptodrosophila latifasciaeformis* males to 4.50 mm in *Zaprionus litos* females [36]. Moreover, the *Drosophila* wing has been extensively studied from the perspectives of morphometrics [37], development [38], and the intersection of the two [39]. This research has revealed the molecular and developmental underpinnings of wing formation as well as the evolutionary processes that have generated both intra-[40, 41] and interspecific variation [42] in wing morphology. The genus thus provides opportunities to understand variation in wing area, body size, and, consequently, wing loading. Nonetheless, most such research has been restricted to *D. melanogaster* and a few other species [43].

Previous work in *D. melanogaster* has been essential to understanding geographic patterns of variation in wing loading. In *D. melanogaster*, high wing:thorax size ratio can serve as proxy for reduced wing loading, and have been proposed to be adaptive for flight at cold temperatures [e.g., 44–48]. Wing:thorax ratio increases with latitude in *D. melanogaster* which has been taken as support for selection favoring large body size, yielding high wing loading, at low temperatures [48].

Here, we expand this body of inquiry to controlled developmental and experimental conditions and comparative methods by quantifying individual-level wing size and body mass to determine the extent of divergence in wing loading among 30 species of drosophilids. We applied comparative phylogenetic methods to determine how much the evolution of mass, wing area, and wing loading are conditioned on phylogeny. We found that while the wing area and body mass show a moderate phylogenetic signal, wing loading does not. Moreover, we find that all the traits are best explained by a model of trait evolution in which trait variation is best explained by changes during speciation events. More conservative analyses indicate no evidence for adaptive peaks in wing loading in *Drosophila*. Our results indicate that the evolution of wing loading in *Drosophila* differs from the evolution of the same trait in birds and bats, and pose the question of whether these differences can be generalized to other insects.

## METHODS

### Stocks and fly collection

We raised flies from 30 drosophilid species (Table S1). We scored isofemale lines whose genomes had been previously sequenced [49]. All flies were raised on standard cornmeal/molasses medium in 200mL bottles provided by Archon Scientific (Durham, NC). Virgin females were collected by lightly gassing recently emerged flies (less than 8 h after eclosion) with CO_2_ and separating females from males. We collected pure-species males and females of each species as virgins within 8 hours of eclosion under CO₂ anesthesia and kept them for three days in single-sex groups of 20 flies in 30 mL vials containing corn meal food. The flies were maintained at 24°C under a 12-hour light/dark cycle for four days. On day 5, flies were anesthetized by placing on an ELISA plate at -20°C for 30-45 minutes (depending on the size of the flies). Table S1 lists all the details of the stocks used in this collection and the available metadata.

### Individual mass

We weighed 10 males and 10 females per species using a Mettler Toledo XPE26 microbalance (Columbus, OH). This device allows for a readability of 1 µg and a repeatability of 0.7 µg. Individuals were placed in the weighing station with metal forceps without making contact with the weighing plate.

### Wings imaging and Image analysis

Immediately after weighing, we removed the wings of each individual with a No. 22 stainless-steel blade permanently mounted on a No. 4 plastic handle (Carolina Biological Supply Company; Burlington, NC) under a Leica dissecting stereoscope (Wetzlar, Germany). One wing per individual was mounted on a pre-clean glass slide (VMR VistaVision^TM^, VWR; cat. no. 16004-422; Radnor, PA) on 20μl of a 1:1 solution of Hoyer’s media and lactic acid. We moved the slides to a baking oven (75°C) for 24 hours to speed clearing and fixing. We captured images of the specimens in JPEG format using an RGB color space using an Olmus scope with an attached camera Gryphax microscope camera (Arktur 8MP SKU: 571025). Wings were photographed at a consistent magnification, and a scale bar slide was photographed at the end of each photography session to facilitate conversion of pixel dimensions to mm for subsequent analyses.

We measured wing area to facilitate calculation of wing loading. We used the ‘Select Subject’ function in Adobe Photoshop [50] to detect the wings within the images. We thresholded the images to create a binary with wings represented by white and the background black. The proximal portion of the wing containing the hinge joint sustained variable amounts of damage during removal from the body, so to standardize the wing area measurements we manually removed that portion of the wing from analysis using Adobe Photoshop. We did this by extending a line from the costal break on the leading edge of the wing to the notch formed between the alula and the main wing membrane on the trailing edge and removing the wing that is proximal to that line from the thresholded image. Figure 1 shows an example of the image analysis routine.

**FIGURE 1.**
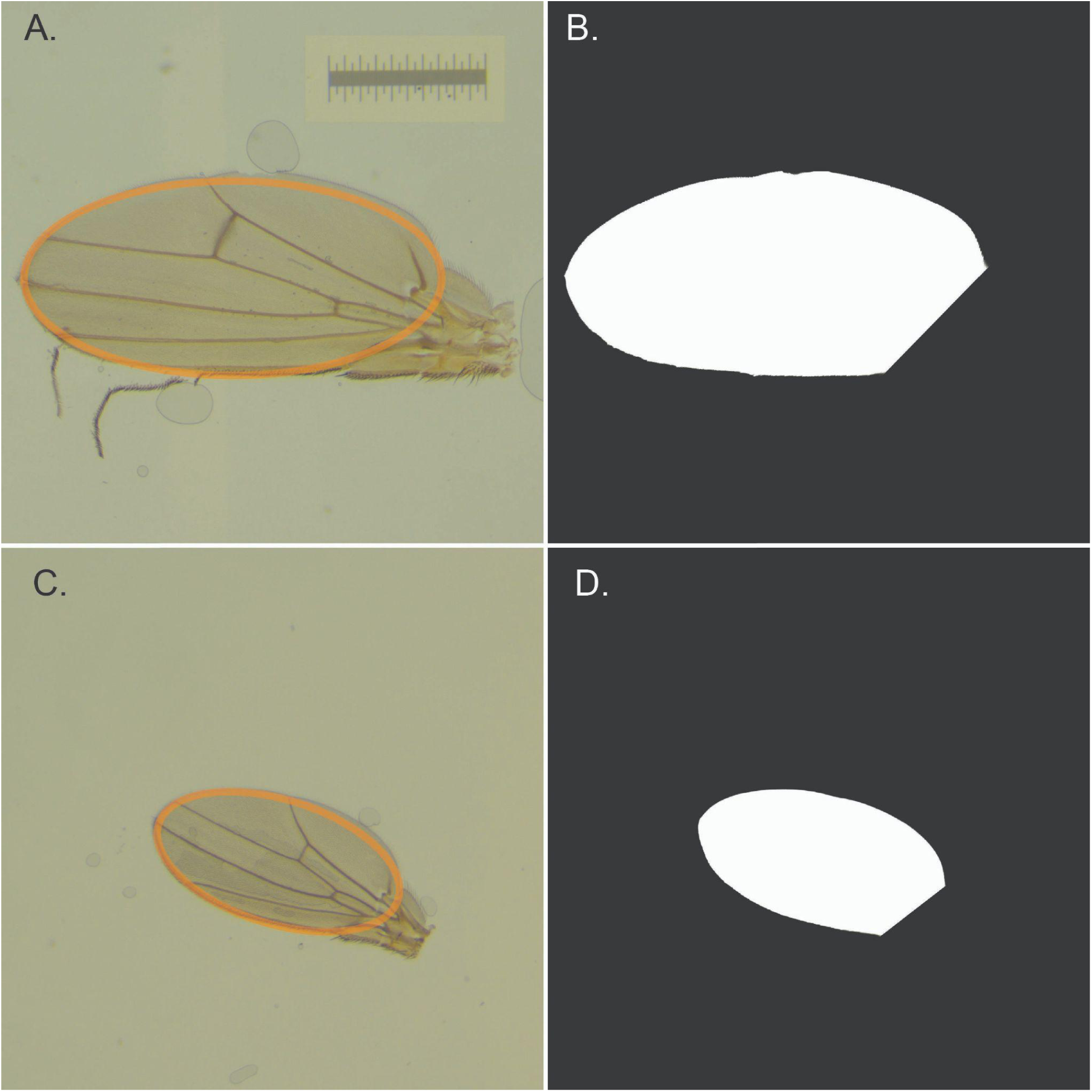
Two examples of wing morphology in the genus *Drosophila*. **A.** Wing from a *D. niveafrons* male under light microscopy. **B.** Same wing when analyzed with the pipeline described in Methods to quantify wing area. **C.** Wing from a *D. nikananu* male. **D.** Same wing when analyzed with the pipeline to quantify wing area. The orange ellipses in A. and C. depict the alternative ellipse method of measuring wing area, also described in Methods. Scale bar = 1 mm, all images presented at the same scale.

We took two area measurements of the white region of each image: 1) A direct count of the number of white pixels (a direct measurement of wing area), and 2) an indirect measurement wherein we calculated an ellipse based on the major (the distance between the alula and the distal tip of the wing) and minor (wing chord) axes of the white image region. We took this approach because several wings were damaged during removal and slide preparation. We found that these two measures were highly correlated (*r* = 0.9998) with less than a 1% difference in the mean areas. We thus decided to use the ellipse measurement method to minimize the influence of wing damage in our calculations of individual wing loading. Measurements were extracted from the thresholded images using the *regionprops* function in Matlab [51]. In addition to the pixel and ellipse area measurements, we also extracted the major and minor axis lengths of the ellipses and calculated the eccentricity of the ellipses. Eccentricity describes the shape of an ellipse and ranges from 0 (a circle) to 1 (a line segment). We photographed a 1mm scale bar at the same magnification to allow us to convert pixel dimensions into real-world measurements (Figure 1).

### Phylogenetic tree

Recent developments have produced a time-calibrated phylogenetic tree using genomewide variation for the genus *Drosophila*. Suvorov et al. [52] and Kim et al. [49] produced a maximum likelihood time-calibrated tree encompassing over 300 species from concatenated genomewide markers. We matched the species in our data set and the tree. The tree was then trimmed using the R function *drop.tip* [library *ape*, 53]. Following pruning, we transformed the tree to an ultrametric topology. We scaled branch lengths by relative time using the *makeChronosCalib* function followed by the *chronos* function [both found in the library *ape*, 53]. The data matrix and the pruned tree were then merged into a comparative dataset using the *comparative.data* function in *phytools* [54].

### Statistical analyses

We used data structure described immediately above to study the mode of evolution of mass, wing area, and wing loading. First, we studied whether drosophilid species differed in their mass and their wing size. We used two types of linear models. First, we used Ordinary Least Squares (OLS) linear models in which wing size and mass were the response variables, and species and sex were the explanatory variables. The models also included the interaction between the species × sex interaction. We used the function *lm* [library *stats*, 55] to fit the linear models. To determine whether the effects and interactions were significant, we used a type-III ANOVA [function *Anova*, R library *car*, 56]. We also used Tukey Honestly Significant Difference post-hoc pairwise comparisons [function *glht*, library *multcomp*, 57] to estimate differences between species pairs, and their significance.

We also used phylogenetically aware models to study whether there were differences in body mass, wing area, and wing loading among species, sexes, and the interaction of species and sex while accounting for the phylogenetic relatedness among species. We used two approaches. First, we performed phylogenetic generalized least squares (PGLS) regression with the functions *gls* (library *nlme* ([58]) and *corPagel* (library ape [53]). For these PGLS models, we used the functions *emtrends* and *emmeans* (library *emmeans* [59]) to perform post-hoc pairwise comparisons. Second, we used randomized residual permutation procedure (RRPP)-based phylogenetic ANOVA with the function *extended.pgls* (library geomorph [60–62]) and 1,000 permutations. We performed two types of phylogenetic ANOVA to complement the lambda-based [63] approach of our PGLS models. In our first phylogenetic ANOVA (RRPP-BM), we used the option *phy* to calculate expected phylogenetic covariances under a Brownian Motion (BM) model of evolution. In our second phylogenetic ANOVA (RRPP-Kappa), we used the option *cov* to to implement the ANOVA with a covariance matrix calculated from our phylogenetic tree rescaled using the function *rescale* (library phytools [54]) with estimates of kappa from the best-fitting evolutionary models for each trait (see Results). We performed post-hoc pairwise comparisons for our phylogenetic ANOVAs with the geomorph function *pairwise* with *test.type* option “stdist”.

### Comparative phylogenetic analyses for mass and wing area

#### Phylogenetic signal

Next, we studied whether body mass, wing area and their ratio, wing loading, followed a pattern of phylogenetic conservatism [i.e., a tendency for related species to share similar mean trait values predictable by their phylogenetic relationships, 64] or if, on the contrary, they had diverged more than expected by their phylogenetic relationships. We used the species tree generated from genome wide markers (described above) to measure phylogenetic signal for each of these three traits using two different complementary metrics. The first one, Pagel’s *λ* [63], estimates how much related species resemble each other more than expected by their genetic relationships; the range of this metric is from 0 (traits evolve randomly along the phylogeny) to 1 (traits are fully predictable by phylogenetic topology). The second metric, Blomberg’s *K* [65], measures how closely related species resemble each other but within the expectation of a Brownian motion model of trait evolution. *K* values lower than 1 suggest greater trait similarity than expected by a Brownian model, which in turn serves as evidence against phylogenetic conservatism. When *K* does not differ significantly from one, the distribution of trait values adheres to the expectations of a Brownian motion. Finally, *K* values larger than 1 indicate that species are more similar in the focal trait than expected. This can suggest (but does not conclusively show) phylogenetic conservatism [66, 67]. We additionally used likelihood ratio tests to compare *λ* values for mass, wing area, and wing loading between males and females.

### Allometry of wing size and wing loading

We evaluated allometry at two different levels: within species and between species. We fit two linear models in which area and wing load, respectively, were expressed as functions of mass, species, and the interaction between these two terms. To account for the potential effect of the interaction, we used the function ANOVA [library *car*, 56] to do a type-3 ANOVA. We evaluated the slope for each species using the function summary [library *stats*, 55].

We also assessed the allometric relationships of wing area and wing loading with respect to mass in a phylogenetic context. To do this, we constructed Phylogenetic Generalized Least Square (PGLS) models with wing area and wing loading, respectively, as functions of mass using the *pgls* function in the R package *caper* [68]. We tested the scaling of the two wing size traits on pooled data (all species and both sexes included), we also parsed the data and assessed sex-specific scaling, and we evaluated whether scaling differed between the two primary drosophilid clades, the subgenus *Sophopora* and *Drosophila*. These models were also constructed using the *pgls* function from the *caper* R package [68]. To highlight the importance of phylogenetic correction in models of allometric scaling relationships, we compared our PGLS results for the wing area vs. body mass relationship to those of an ordinary least-squares model, without phylogenetic corrections.

### Evolutionary models of trait evolution for mass, wing area, and wing loading

Next, we investigated whether wing area, individual mass, and wing loading were each described by a best-fitting model of evolution along the phylogeny. We used the R function *fitcontinuous* [library *geiger*, 69, 70] to quantify the relative support for each of eight evolutionary models listed in Table S2. The models represented a range between no relationship between trait evolution and the phylogeny (white noise) to variations in the *tempo* and mode of evolution with including random evolution (Brownian motion), constrained random evolution [Ornstein-Uhlenbeck, 71], and a variety of branch length transformations that simulate changes in evolutionary rate across the tree (early-burst, kappa, delta, and rate-trend, [63, 69, 70, 72, 73]). Table S2 lists the description and parameters for each model. To compare among models, we estimated their sample-size corrected Akaike Information Criterion (AICc) values and compared their relative support using Akaike weights [wAIC, 74].

### Ancestral character state reconstructions

We performed ancestral character state reconstruction [75] to describe the evolutionary history of *Drosophila* traits. We used the R function *anc.ML* function in the R package phytools [54] to apply a Brownian motion model to estimate the change in body mass, wing size, and wing loading on the pruned *Drosophila* phylogeny. We extracted the estimated values for each node within the tree along with their respective 95% confidence intervals. To assess whether the pattern of trait evolution was gradualistic or if any of the branching events in the tree corresponded with rapid morphological divergence, we calculated the overlap between confidence intervals of the parent nodes and each of their child nodes. Overlapping confidence intervals would suggest gradual evolution while non-overlap would suggest an event of rapid divergence.

### Evolutionary rate analyses and evolutionary models of trait evolution for wing loading

We investigated the pace of morphological evolution in drosophilid wings. We calculated the evolutionary rate parameter (*σ*^2^ - the rate of change in trait mean under a Brownian motion process) using the *fitcontinuous* function as described above. We used the wAIC values to compare support for the evolutionary rate estimates generated by each of the evolutionary models. We also modeled evolutionary rate patterns to assess whether trait evolution were consistent across the *Drosophila* phylogeny, or whether they varied throughout the history of the lineage. To do this, we conducted maximum-likelihood models of evolutionary rate of body mass and wing area with the *multirateBM* function in R [library *phytools*, 54]. These models use penalized likelihood to fit a flexible Brownian motion rate to the tree [76], essentially estimating ancestral shifts in the pace of evolution over time. To facilitate comparison of the resulting patterns between the traits, body mass, and wing area, we rescaled the *σ*^2^ output from each *multirateBM* model to a range of 0-1. We then subtracted the rates for wing area from those for body mass, creating an index of relative change ranging from -1 to 1. At an index value of 0, both traits are evolving at their mean rates; values greater than 0 show relatively slow evolution of wing area relative to body mass, and values less than 0 show relatively rapid evolution of wing area relative to mass.

## RESULTS

### Mass and wing area differ among *Drosophila* species and sexes

#### Wing area

The range of wing area was large, and the ratio between the largest and smallest wings we scored was 4.15. *Drosophila niveifrons* had the wings with the largest area (*n*=20, mean = 2.706 mm^2^, sd=0.301 mm^2^); *D. nikananu* was the species with the smallest mean wing area (*n*=20, mean = 0.928 mm^2^, sd=0.086 mm^2^; Figure 2). We estimated the extent of phenotypic differentiation in wing area across species and sexes. Both the OLS species effect (F_29,537_ = 338.883, P < 1 × 10^-15^) and sex effect (F_1,537_ = 439.126, P < 1 × 10^-15^) were strong. The interaction between sex and species was significant (F_29, 537_ = 7.013, P < 1 × 10^-15^), and post-hoc pairwise tests of sexual dimorphism (female-male means) revealed significant differences for 25 of 30 species (Figure 4, Tables S3-S4). Post-hoc tests also confirmed significant differences in the majority of pairwise tests of species differences (Figure S1, Table S5).

**FIGURE 2.**
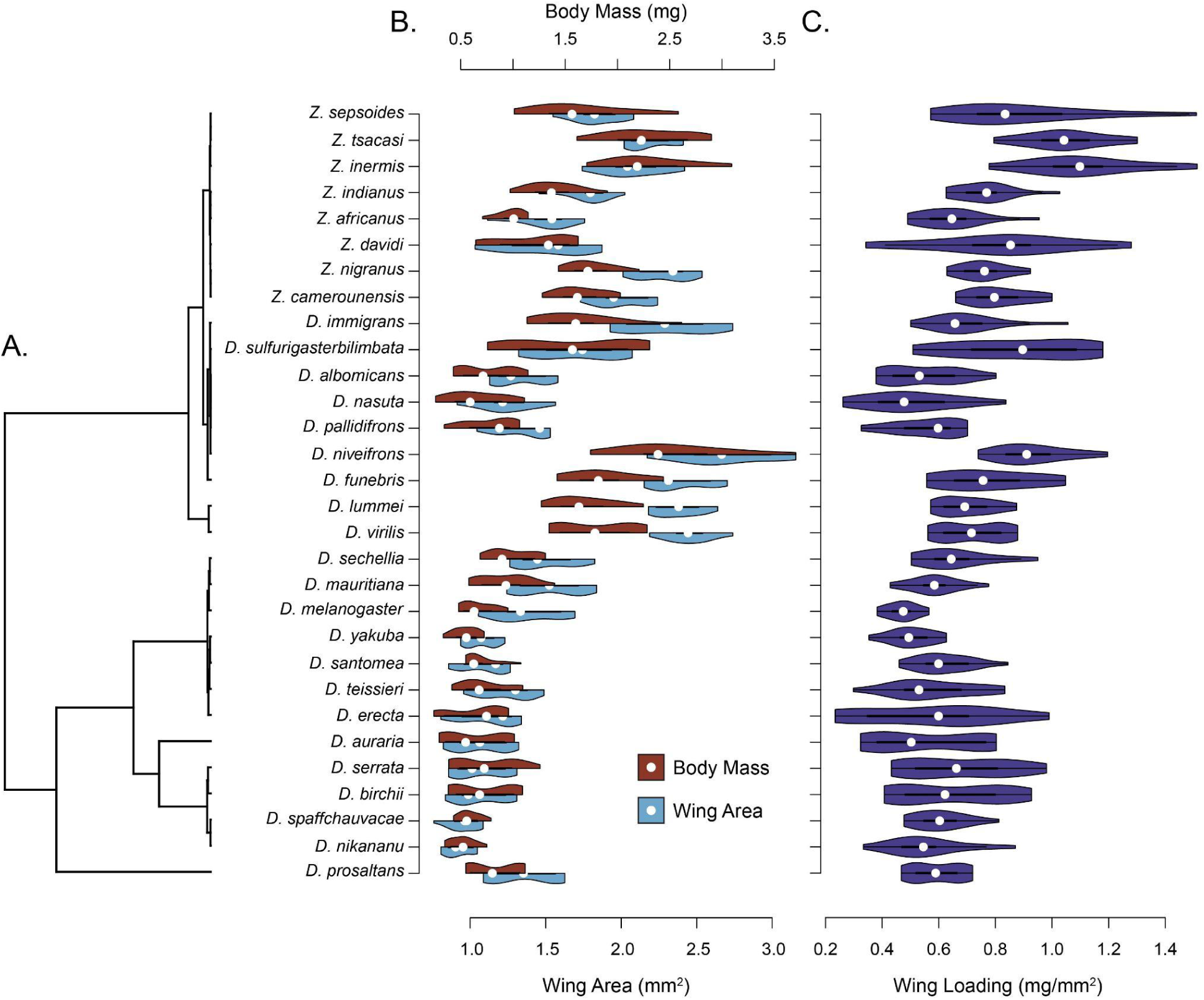
Species tree (A.) and phylogenetic distribution of body mass, wing size (B.), and wing loading (C.) in 30 drosophilid species. Panel A. shows the ultrametric drosophilid tree upon which our comparative analyses are based. Violins in Panels B. and C. show the distributions of each of the measured traits: red upper violin halves in A. show body mass paired with wing area (blue lower half violins), and violet violins in C. show wing loading.

**FIGURE 3.**
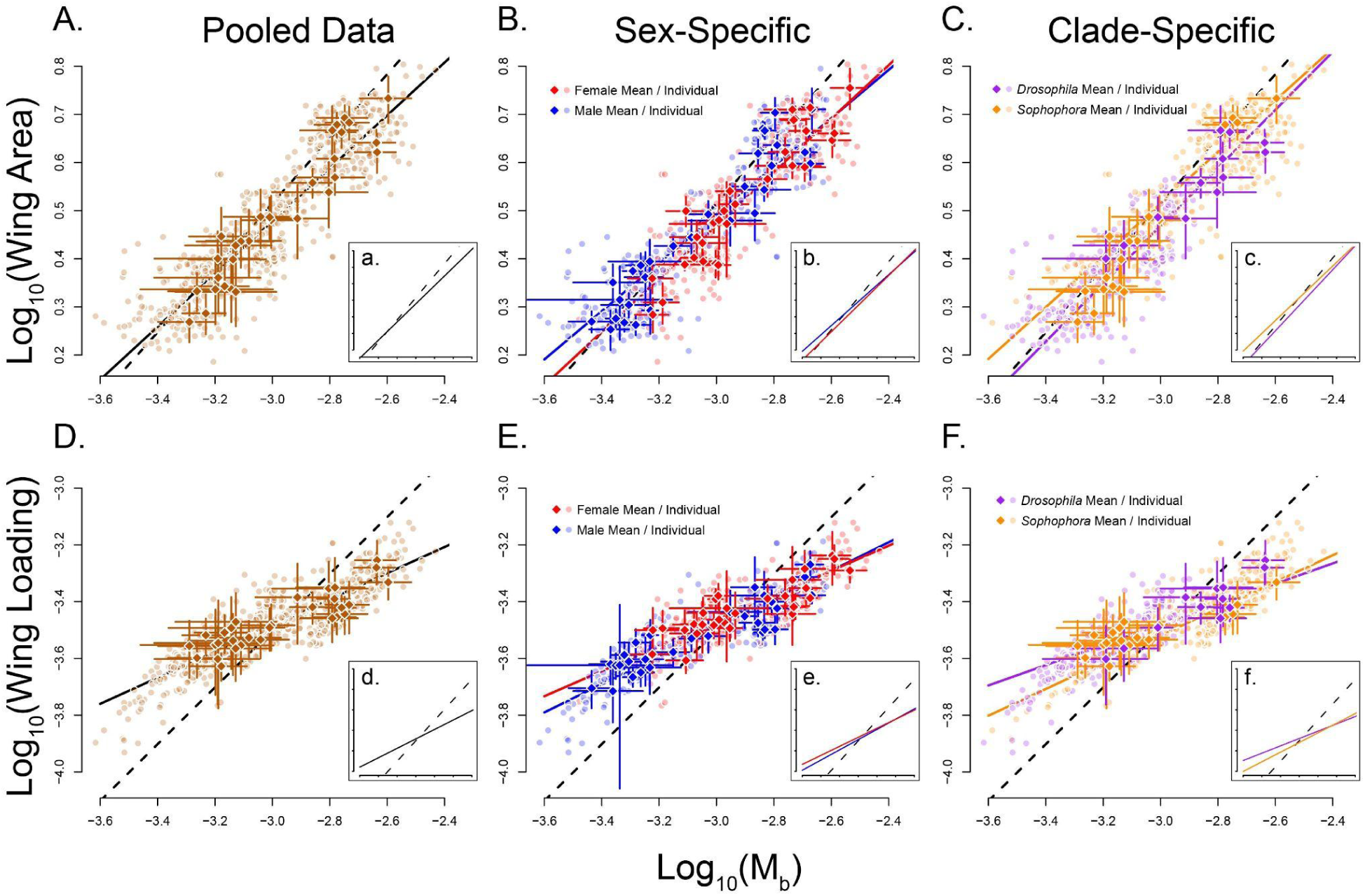
Wing area and wing loading scale positively with respect to body mass in *Drosophila*, but fall short of isometric scaling. We analyzed scaling of wing area (A.-C.) and wing loading (D.-F.). In three partitioning schemes the complete pooled dataset (A., D.), segregated by sex (B., E.), and by the two primary *Drosophila* subgenera (*Sophophora* and *Drosophila*; C., F.). Translucent points show individual measurements, and solid-color diamonds show species means with ±1 sd error bars. Solid lines show PGLS regression models compared to the isometric expectations, shown by dashed lines. Insets (a.-f.) offer comparison of regression slopes with isometry with data points removed for clarity.

**FIGURE 4.**
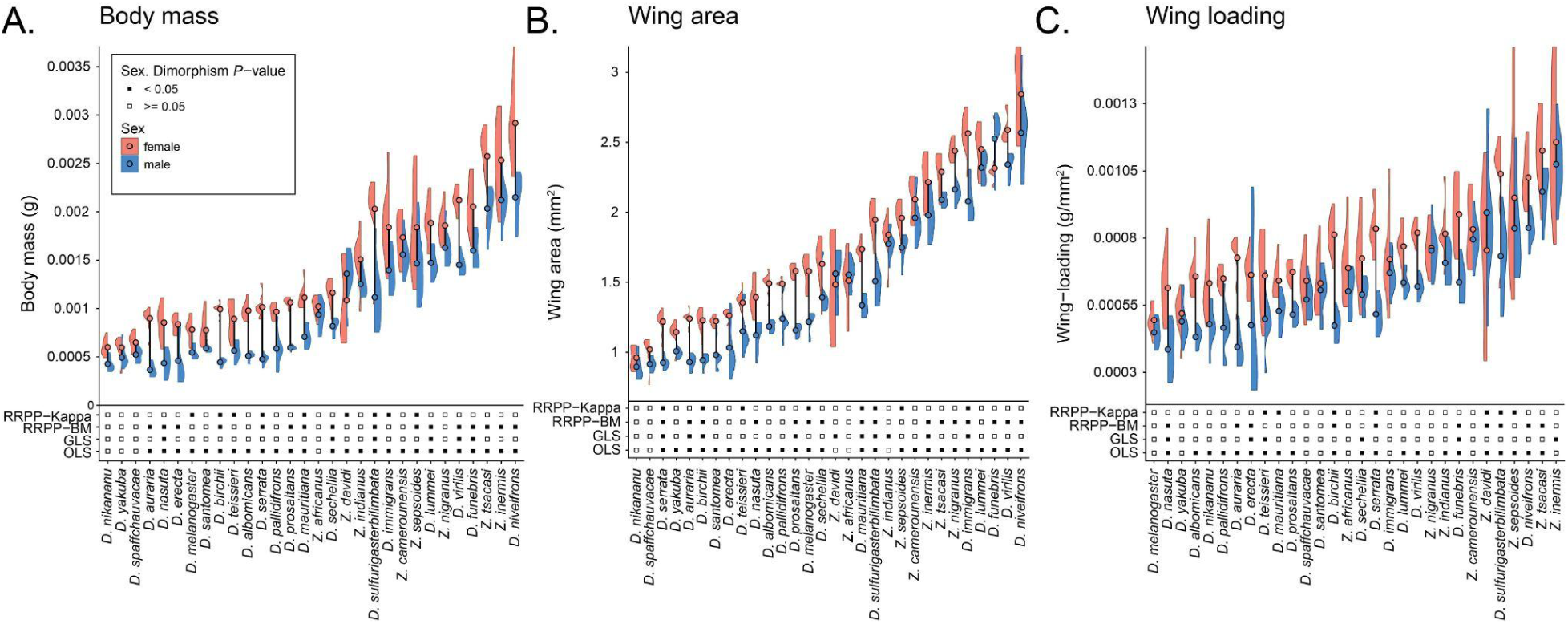
Sex differences in body mass, wing area, and wing loading among 30 *Drosophila* species. Species were ordinated in ascending order and arranged to show the lowest mean of each trait, divided by sex, on the left and highest on the right. In each panel, species are on the x-axis and significance for model-specific tests of pairwise differences between sexes in each species is indicated immediately above the x-axis. Y-axes above the model information in each panel correspond to data for body mass (A.), wing area (B.), and wing loading (C.), with separate female (red) and male (blue) distributions and means (like-colored points) plotted for each species. Thick black lines connect male and female means for each species.

Our phylogenetic models produced varied results. All three model terms (Species, Sex, and their interaction) were highly significant for both PGLS (Species: F_29,537_ = 9.082, P < 1 × 10^-4^; Sex: F_1,537_ = 21.110, P < 1 × 10^-4^; Interaction: F_29,537_ = 2.673, P < 1 × 10^-14^) and RRPP-BM (Species: F_29,537_ = 25.544, P < 1 × 10^-3^; Sex: F_1,537_ = 435.769, P < 1 × 10^-3^; Interaction: F_29,537_ = 7.080, P < 1 × 10^-3^), but only Sex and its interaction with Species were significant for RRPP-Kappa (Species: F_29,537_ = 0.417, P = 0.998; Sex: F_1,537_ = 7.057, P < 1 × 10^-3^; Interaction: F_29,537_ = 4.330, P < 1 × 10^-3^). Post-hoc tests of sexual dimorphism varied, with significant differences for 10, 16, and 8 of 30 species for PGLS, RRPP-BM, and RRPP-Kappa, respectively (Figure 4, Tables S6-S7, S9-S12). Similarly, post-hoc comparisons of interspecific differences varied, but for all models, the number of significant pairwise tests was much lower than for OLS (and was 0 for RRPP-Kappa) (Figures S2-S4; Tables S6, S8-S12). These lower proportions of significant tests for all phylogenetic models relative to OLS suggest these analyses may be underpowered with our sample size of 20 individuals (10 male and 10 female) per species. A non-mutually exclusive alternative is that phylogenetic history may account for most of the species-level variance in mean wing area.

#### Mass

Next, we studied whether mass differed among *Drosophila* species. The results follow a pattern similar to that observed in our analyses of wing area. We find that the species effect is significant (F_29,537_ = 182.319, P < 1 × 10^-15^) which is not surprising as the range of body mass is nearly fivefold (*D. niveifrons*: *n*=20, mean = 2.534 × 10^-3^ g, sd= 5.150 × 10^-4^ g vs. *D. nikananu*: *n*=20, mean= 5.138 × 10^-4^ g, sd=1.1422 × 10^-4^ g). The sex effect was also significant (F_1,537_ = 521.623, P < 1 × 10^-15^), as was its interaction with species (F_29,537_=6.574, P < 1 × 10^-15^). Post-hoc tests of sexual dimorphism were significant for 26 of 30 species (Figure 4, Tables S13-S14), and most post-hoc pairwise comparisons among species were significant, as well (Figure S5, Table S15).

As was the case with wing area, our phylogenetic models produced varied results for body mass. PGLS estimates of Species (F_29, 537_ = 7.263, P < 1 × 10^-4^) and Sex (F_1, 537_ = 44.435, P < 1 × 10^-4^) were significant, but their interaction was marginal (F_29, 537_ = 1.401, P = 0.079). All three model terms were significant for RRPP-BM (Species: F_29, 537_ = 21.061, P < 1 × 10^-3^; Sex: F_1, 537_ = 521.123, P < 1 × 10^-3^; Interaction: F_29, 537_ = 6.548, P < 1 × 10^-3^), but, as was the case for wing area, the species term for RRPP-Kappa was not significant (Species: F_29, 537_ = 0.286, P = 1.000; Sex: F_1, 537_ = 361.548, P < 1 × 10^-3^; Interaction: F_29, 537_ = 8.288, P < 1 × 10^-3^). Post-hoc tests of sexual dimorphism varied, with significant differences for 6, 16, and 9 of 30 species for PGLS, RRPP-BM, and RRPP-Kappa, respectively (Figure 4, Tables S16-S17, S19-S22). Similarly, post-hoc comparisons of interspecific differences varied, but for all models, the number of significant pairwise tests was much lower than for OLS (Figures S6-S8; Tables S18, S20, S22).

### Allometric relationships between wing size and mass

We evaluated whether the wing area scaled with body size, using mass as a proxy of the latter. We found a strong correlation between the two metrics and that this correlation deviated from the expectation of isometric scaling (i.e., slope equal to 0.66 to reflect the expected area vs volume relationship). Wing area scaled relative to mass with a scaling factor of 0.55 ± 0.048 in our PGLS model (*F*_1,28_ = 130.3, *P* = 4.820 x 10^-10^, adjusted *R*^2^ = 0.82), which was significantly lower than 0.66. This result also diverges from the results of a non-phylogenetically controlled ordinary least-squares regression (OLS) model, in which we found a scaling factor of 0.61 ± 0.039 (*F*_1,28_ = 253.1, *P* = 1.483 x 10^-15^, adjusted *R*^2^ = 0.90) that was was not statistically distinct from the isometric null (Wald *χ*^2^ test, P = 0.18). The phylogenetic relatedness of the sampled taxa means that OLS is not an appropriate test for these data [77], and our results highlight the importance of including phylogenetic correction in models of interspecific allometric scaling. All other allometric models that we present here are phylogenetically aware PGLS models.

We also explored whether wing size scaled differently between sexes and the two primary clades of *Drosophila*. Wing area scaling differed between males and females; males had a scaling factor of 0.50 ± 0.046 (*F*_1,28_ = 115.0, *P* = 2.017 x 10^-11^, adjusted *R*^2^ = 0.80) and females had a scaling factor of 0.55 ± 0.051 (*F*_1,28_ = 118.6, *P* = 1.413 x 10^-11^, adjusted *R*^2^ = 0.80). Wing loading in the pooled dataset (including males and females in the same model) scaled with a factor of 0.46 ± 0.048 (*F*_1,28_ = 91.77, *P* = 2.463 x 10^-10^, adjusted *R*^2^ = 0.76). As with wing area, wing loading scaling factors differed between males and females (0.50 ± 0.047 and 0.44 ± 0.051, respectively). Scaling factors also differed between the two *Drosophila* clades. The *Drosophila* subgenus had a wing area scaling factor of 0.54 ± 0.06, versus 0.60 ± 0.12 for the *Sophophora* subgenus. Wing loading in the *Drosophila* subgenus had a scaling factor of 0.47 ± 0.56, compared to 0.36 ± 0.12 in the *Sophophora* subgenus.

### Wing loading in the *Drosophila* genus

Our survey measured wing area and mass for the same individuals which allowed us to quantify wing loading as a direct measurement for each individual. We found that the range of WL values is more than twofold with *Z. inermis* having the largest value (*n* = 20, mean = 1.114 × 10^-3^, sd = 1.899×10^-4^ g/mm^2^) and *D. melanogaster* having the lowest (*n* = 20, 4.713 × 10^-4^, sd = 5.267×10^-5^ g/mm^2^). Note that *D. melanogaster* is not the species with the lowest wing area or the highest mass, so the low wing loading is not the result of a body dimension outlier. As expected, OLS revealed significant differences among species (F_29,537_ = 37.347, P < 1 × 10^-15^), and both Sex (F_1,537_ = 220.195, P < 1 × 10^-15^) and its interaction with species (F_29,537_ = 5.078, P < 1 × 10^-10^) were also significant for OLS. Of the 30 evaluated species, post-hoc pairwise comparisons detected significant sexual dimorphism in 21 species (Figure 4, Tables S23-S24). Of these 21 dimorphic species, females showed higher wing loading than males for 20 species, while one species, *Z. davidi*, exhibited higher wing loading for males. Overall, these results indicate the existence of sexual dimorphism in *Drosophila* wing loading and that, where it occurs, females tend to have higher wing loading than males. Fewer than half of all OLS post-hoc pairwise species comparisons were significant (Figure S9, Table S25).

As was the case with wing area and body mass, our phylogenetic models produced varied results for wing loading. PGLS estimates of Sex (F_1, 537_ = 30.006, P < 1 × 10^-4^), Species (F_29, 537_ = 2.653, P < 1 × 10^-4^) and their interaction were significant (F_29, 537_ = 1.696, P = 0.0139). All three model terms were also significant for RRPP-BM, (Species: F_29, 537_ = 3.783, P < 1 × 10^-3^; Sex: F_1, 537_ = 220.393, P < 1 × 10^-3^; Interaction: F_29, 537_ = 4.191, P < 1 × 10^-3^), but, as was the case for wing area and body mass, the species term for RRPP-Kappa was not significant (Species: F_29, 537_ = 0.063, P = 1.000; Sex: F_1, 537_ = 142.348, P < 1 × 10^-3^; Interaction: F_29, 537_ = 5.477, P < 1 × 10^-3^). Post-hoc tests of sexual dimorphism varied, with significant differences for 6, 10, and 7 of 30 species for PGLS, RRPP-BM, and RRPP-Kappa, respectively (Figure 4, Tables S26-S27, S29-S32). Similarly, post-hoc comparisons of interspecific differences varied, but for all models, the number of significant pairwise tests was much lower than for OLS (Figures S10-S12; Tables S28, S30, S32).

### Phylogenetic signal and evolutionary models

Next, we evaluated whether mass, wing area, and wing loading showed a significant phylogenetic signal, which in turn measures how phylogenetic history conditions trait divergence. Both mass and wing size showed a moderate value of Pagels’s *λ* (Table 1) that differed significantly from 0 (P < 1.0 × 10^-4^), but in neither case was close to 1. These results indicate that the species-level means of these two traits are influenced, but not completely conditioned, by phylogenetic history.

**TABLE 1.** Metrics of phylogenetic signal in body size, wing area, and wing loading for drosophilid species.

| Signal Metric | Mass<br>($P=0$, $P=1$) | Area<br>($P=0$, $P=1$) | WL<br>($P=0$, $P=1$) |
| --- | --- | --- | --- |
| Pagel's $\lambda$ | 0.400<br>($P=0.00050$, $P=0.00$) | 0.511<br>($P=0.00012$, $P=0.00$) | 0.296<br>($P=0.01310$, $P=0.00$) |
| Blomberg's $K$ | $2.963 \times 10^{-3}$<br>($P=0.064$, $P=0.0010$) | $4.147 \times 10^{-3}$<br>($P=0.014$, $P=0.0010$) | $3.092 \times 10^{-3}$<br>($P=0.073$, $P=0.0010$) |

**TABLE 2.** Fit of eight models of trait evolution to patterns of variation in wing area, body mass, and wing loading (WL) in drosophilids. BM: Brownian motion, OU: Ornstein–Uhlenbeck, EB: Early-burst.

|  | Area |  | Mass |  | WL |  |
| --- | --- | --- | --- | --- | --- | --- |
|  | AIC<sub>Area</sub> | wAIC<sub>Area</sub> | AIC<sub>Mass</sub> | wAIC<sub>Mass</sub> | AIC<sub>WL</sub> | wAIC<sub>WL</sub> |
| BM | 89.976 | $1.363 \times 10^{-3}$ | -305.789 | $4.95 \times 10^{-16}$ | -384.24 | $5.853 \times 10^{-15}$ |
| OU | 47.723 | $2.039 \times 10^{-5}$ | -358.256 | $1.225 \times 10^{-4}$ | -438.379 | $3.338 \times 10^{-3}$ |
| Lambda | 37.640 | $3.155 \times 10^{-3}$ | -366.267 | $6.727 \times 10^{-3}$ | -439.038 | $4.641 \times 10^{-3}$ |
| Kappa | **26.129** | **0.997** | **-376.256** | **0.993** | **-449.753** | **0.985** |
| Delta | 47.069 | $2.829 \times 10^{-5}$ | -358.667 | $1.505 \times 10^{-4}$ | -439.714 | $6.507 \times 10^{-3}$ |
| EB | 92.455 | $2.829 \times 10^{-5}$ | -303.311 | $1.436 \times 10^{-16}$ | -381.761 | $1.695 \times 10^{-15}$ |
| White noise | 49.898 | $6.877 \times 10^{-6}$ | -356.64 | $5.462 \times 10^{-5}$ | -435.361 | $7.382 \times 10^{-4}$ |
| Trend | 89.70879 | $1.558 \times 10^{-14}$ | -306.063 | $5.685 \times 10^{-16}$ | -384.514 | $6.712 \times 10^{-15}$ |

We also calculated Blomberg’s *K*, a metric that reflects the magnitude of the phylogenetic signal under the expectation of a Brownian Model of trait evolution, for mass and area. *K* values for both traits were low and did not differ from zero. We thus compared the fit of our data to seven models of trait evolution. These models represented a variety of trait evolutionary modes. We found that of the seven tested evolutionary models, the best fit for both traits was the kappa (speciational) model of trait evolution. This model of trait evolution modifies branch lengths in a phylogeny to test whether trait evolution is proportional to the amount of evolutionary time represented by those branches. A more streamlined comparison that included only the White Noise (WN), Brownian Motion (BM), and Ornstein–Uhlenbeck (OU) models indicated that the OU model was the most likely to explain trait evolution in both body size and wing area. In both cases, AIC_w_ was at least twice as large for the OU model as for the BM model, indicating the existence of stabilizing selection and optimal trait values in wing area (Table S3).

We also estimated metrics of phylogenetic signal for the ratio of wing area and mass, wing loading. We found a more modest signal in Pagel’s *λ* than in mass or wing area, and also found a non-significant Blomberg’s *K* (Table 1, Figure 5). Similar to the results from mass and wing area, the best evolutionary model to explain the distribution of wing loading in *Drosophila* was the kappa model (Table 1). When we restrict these analyses to the three basic modes (WN, OU, and BM), wing loading shows overwhelming support for an OU evolutionary model (Table S33). These results suggest that wing loading has a more labile signal than both size and wing area when interpreted in isolation but that the model of evolution is similar to its two components. Overall, none of the traits (mass, wing area, and wing loading) conform to the expectations of a random model of evolution dominated by drift (i.e., BM) and instead the phylogenetic signal in the traits is best explained by a model of divergence at the splitting instances in the evolution of *Drosophila*, and potentially to trait optimal values. Phylogenetic signal did not differ between sexes in any of the measured traits (likelihood ratio tests, all *P* > 0.5)

**FIGURE 5.**
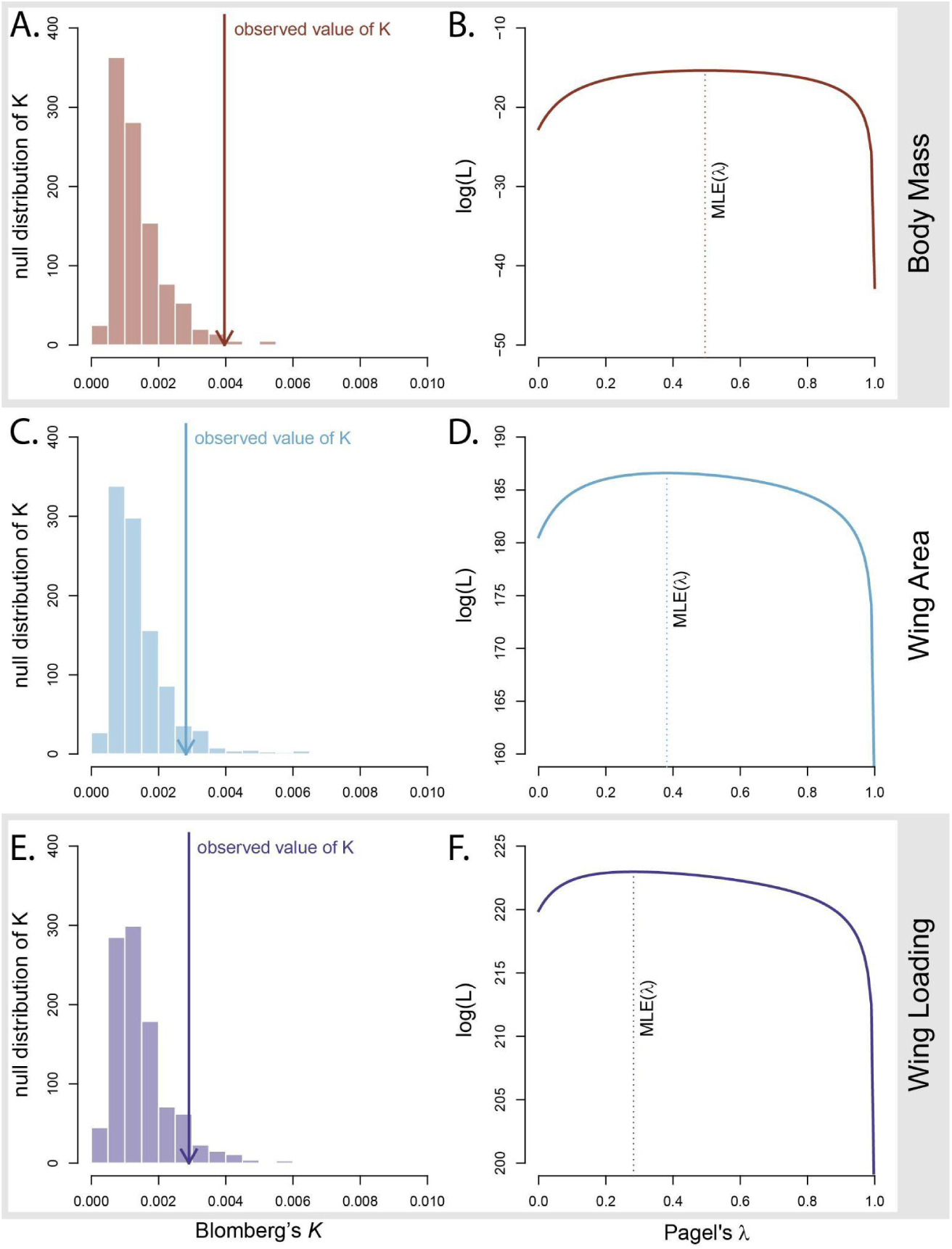
Phylogenetic signal in wing loading in *Drosophila*. Pagel’s *λ* and Blomberg’s *K* are two metrics of phylogenetic signal (described in the text). **A**. Pagel’s *λ* value for female body mass in *Drosophila*. Estimated likelihood of Pagel’s *λ* values. The dotted line marks the maximum likelihood estimate (MLE) of *λ*. **B**. Blomberg’s *K* for female body mass in *Drosophila*. The histogram shows the distribution of simulated values of Blomberg’s *K*; the blue arrow shows the observed value of Blomberg’s *K*. **C.** Pagel’s *λ* value for wing area. **D.** Blomberg’s *K* for wing area in *Drosophila*. **E.** Pagel’s *λ* value for wing loading. **F.** Blomberg’s *K* for wing loading.

We also explored the evolutionary history of body size, wing size, and wing loading across the *Drosophila* phylogeny. The two primary *Drosophila* clades have diverged in all three traits; members of the *Zaprionus* group are, on average, larger than species in the *Sophophora* group in all three traits. We used ancestral character state reconstruction to ask whether these trait divergences occurred rapidly at branching events within the tree or more gradually. We found that the confidence intervals of parent nodes within the tree generally overlapped those of their daughter nodes, indicating gradualistic evolution rather than abrupt divergence events. The lone exception to this pattern occurred at the branch between *D. pallidifrons* and the sister species *D. nasuta* / *D. albomicans*, the latter two and the node between them having larger body mass and greater wing loading than *D. pallidifrons* and their parent node (Figure 6). Despite this pattern of generally incremental evolution, the ratio of wing size to body size was not consistent among taxa (See Figure 6). In some cases, wing area was relatively large relative to body size, as in the *D. lummei* / *D. virilis*, *D. sechellia* / *D. mauritiana*, and *Z. nigranus* / *Z. camerounensis* clades. This mismatch led to relatively low wing loading in these clades (3.55 ± 0.50×10-4, 3.10 ± 0.55×10^-4^, and 3.93 ± 0.47×10^-4^ g/mm^2^, respectively). Conversely, however, the clade containing *Z. sepsoides*, *Z. tsacasi* and *Z. inermis* had relatively small wings when compared to their body size, leading to relatively high wing loading (4.46 ± 1.21×10^-4^, 5.25 ± 0.07×10^-4^, 5.57 ± 0.10×10^-4^ g/mm^2^, respectively; 5.09 ± 1.06×10^-4^ overall). *Drosophila niveifrons* is an exceptionally large fly (2.39 ± 0.52×10^-3^ g) in which we see high wing loading (4.66 ± 0.06×10^-4^ g/mm^2^). This species reflects the functional outcome of isometric scaling and dimensional mismatch - because body mass scales as a function of length^3^ while wing area scales as a function of length^2^, the inevitable outcome of morphological isometry is that large taxa will be constrained by higher wing loading.

**FIGURE 6.**
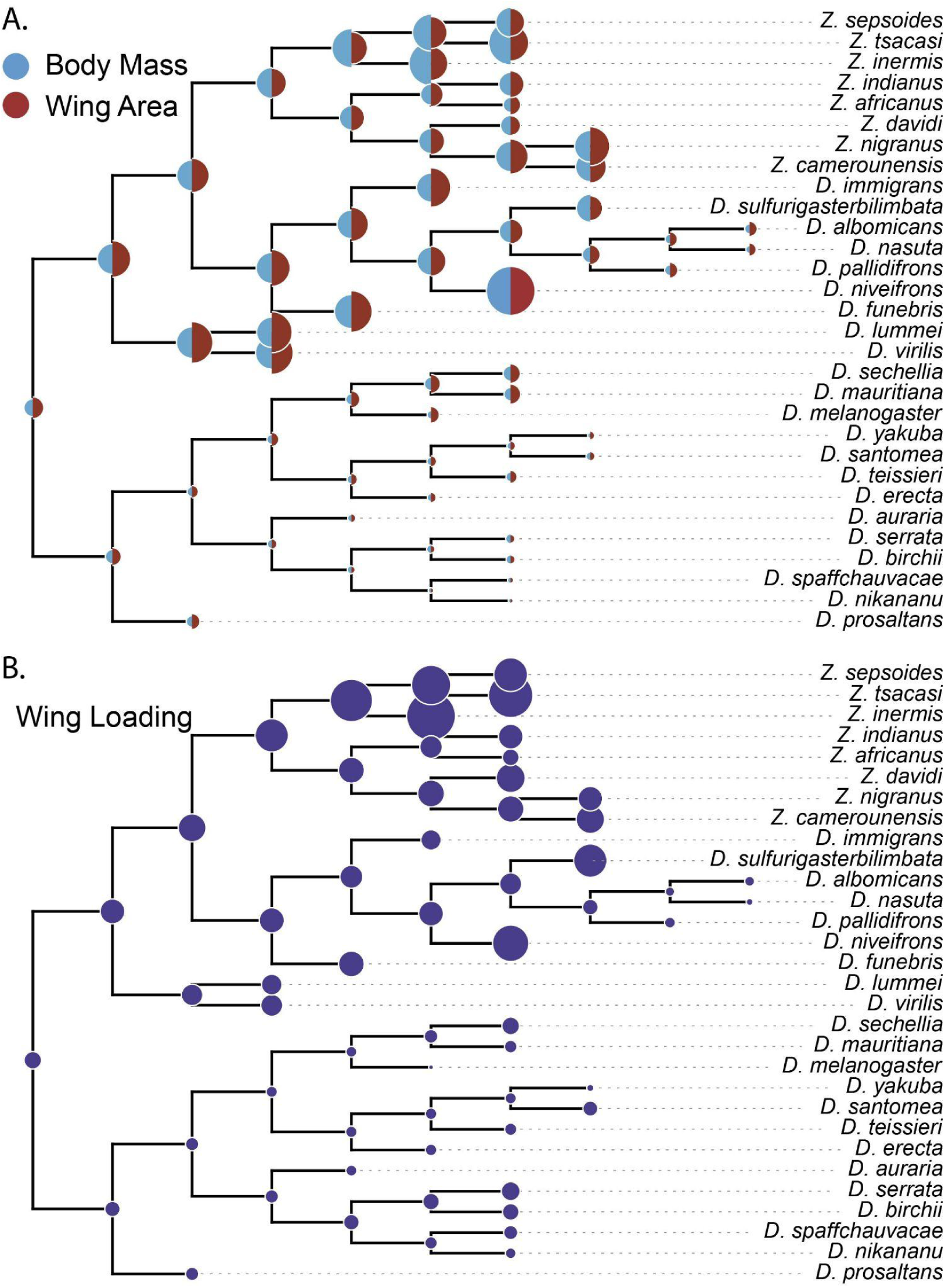
Ancestral state reconstruction of body mass and wing area (A.), and their ratio, wing loading (B.). Marker size was scaled relative to the size range of each trait. Blue semicircles (left half of markers in Panel A.) depict species mean values of body mass at branch tips, and the results of our ancestral character state reconstruction at the interior tree nodes. Red semicircles (right half of makers in A.) depict wing area. Violet markers in B. show species means and reconstructed ancestral character state values for wing loading. Tree branch lengths were all set equal to 1 to display branching events; true tree topology is shown in Figure 2.

We additionally found that evolutionary rates of wing area and body mass generally follow similar patterns (Figure 7). Our multi-rate analysis showed that evolutionary rates varied almost directly between these traits across much of the tree, with a few exceptions. The clade of *Z. sepsoides*, *Z. tsacasi* and *Z. inermis*, in which we saw large body size paired with relatively small wings, also showed relatively slow evolution of wing size when compared to the pace of body size evolution. By contrast, the remainder of the *Zaprionus* showed relatively rapid change in wing size relative to body size.

**FIGURE 7.**
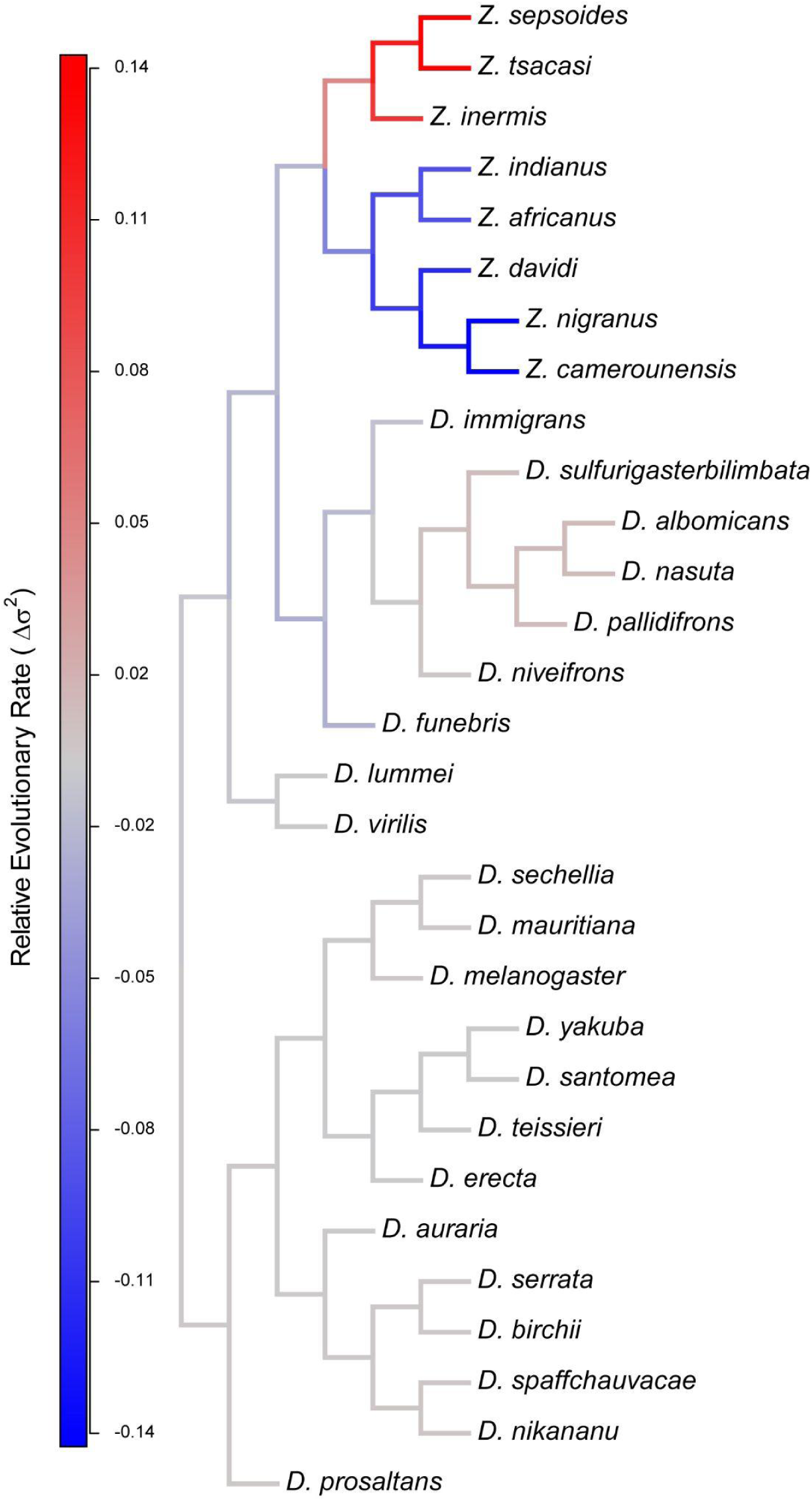
Divergence in ancestral character state reconstruction of evolutionary rate of wing area and body mass. Tree branch lengths were all set equal to 1 as in Figure 6. The color on the branches represent a comparison of evolutionary rate estimates (described in Methods). When branches are gray, body mass and wing area are evolving at their mean rates. Red branches show faster evolutionary rates in body size compared to wing area, and blue branches highlight the reverse: relatively fast evolution of wing area compared to body mass.

## DISCUSSION

In this report, we catalogued the extent of inter and intraspecific variation in wing loading for 30 species of *Drosophila* when all were raised under the same conditions. We find large differences in mass and wing area across species, and in some species, between sexes. We used this survey to estimate wing loading at the individual level for males and females of each species. We also report that, as expected, there is a positive correlation between wing area and mass. This relationship has a slope lower than expected by pure morphological isometry and thus constitutes evidence of negative allometry in the wing area in *Drosophila.* The within-species relationships are not always positive and are sometimes contingent on sex, indicating the existence of sexual dimorphism in scaling of wing area. Next, we used comparative methods to determine how much trait evolution is conditioned by phylogenetic history; we found a moderate phylogenetic signal in mass, wing area, and less so, wing loading, which indicates that this latter trait is more evolutionary labile than expected by just phylogenetic relationships between species. We did not find evidence that phylogenetic signal varies between sexes, despite the dimorphism in some species. In addition to our reports of wing loading and wing area, we report individual sex-specific mass distributions for 30 species. The data we report here will be of use for further comparative physiology analyses. We discuss the findings of sex-differences in wing loading, negative allometry, and phylogenetic signal in the next paragraphs.

Wing loading determines flight speed [78], maneuverability [79], and even diving performance in aquatic birds [80]. Species or individuals with low wing loading can generate more lift at lower speeds, facilitating takeoff, tight turning, and sustained soaring in weak updrafts, whereas high wing loading favors faster flight and improved performance in strong winds but requires higher takeoff speeds and greater power output [2, 81]. Wing loading is often associated with habitat use [82–84], and dispersal [32, 85, 86]. Our results show that wing loading in some drosophilids can be twice the value of other species, indicating different flight strategies among species. The evolution and functional implications of species-level differences in wing loading among *Drosophila* species will require a better understanding of each species’ functional ecology factoring aspects like migratory capacity, and territoriality.

Our results indicate that the evolution of wing loading in *Drosophila* is different from the patterns of evolution in vertebrates. The phylogenetic signal of wing loading is much higher in horseshoe bats (*λ*=0.986, *K*=1.327; [87]) and waterbirds (*λ*=0.827, [88]) than we observed in *Drosophila*. The precise reasons for these differences are unknown but might be related to the distances different *Drosophila* species cover during the lifespan. Experiments with *D. melanogaster* show that some individuals can disperse up to 12 km in a single flight in still air but could cover more distance in a moderate wind [89–92]. These distances may be species-specific, following the differences in flight performance that our present results suggest. Species from the *yakuba* species group cover distances in the hundreds of meters in a day (less if they are hybrids, [93]). A similar pattern is observed in *D. falleni*, a mushroom feeder [94], and *D. suzukii* [95], but see [96]. Evaluating whether the evolution of wing loading differs between insects, plants with dispersing seeds, and flying vertebrates will need a phylogenetically informed perspective. While some efforts have quantified the extent of wing loading in stick insects [97], beetles [32, 98, 99], and wasps [100], none of these efforts has leveraged the power of comparative phylogenetics. Datasets for wing loading already exist (e.g. [34]), which, combined with the emergence of large-scale phylogenomics for these groups, will allow for formal tests across taxa.

Our results indicate that in *Drosophila,* there is no universal sexual dimorphism in wing loading; but when species show sex differences, females usually have higher wing loading. Other insect species also show results consistent with this sex × species interaction. Female Colorado potato beetles (*Leptinotarsa decemlineata*) show higher wing loading than males [101]. Similarly, in some *Cicada* species—but not all—, females have larger wing loading values than males [102], which in turn has been interpreted as selection on flight maneuverability, suggesting these traits are important for reproductive success. Female butterflies *Euphydryas phaeton* show twice the wing loading as males during most of their life (wing loading becomes equivalent after wings are worn), which might be correlated with the habitat choice of the two sexes. Females usually dwell in sheltered and warmer microenvironments while males spend more time basking [103]. An even more extreme case is found in the cricket *Phymateus morbillosus*, in which females have wing loading so high that they are flightless [104], while males, with lower wing loading, are capable of flight. On the other hand, whitetail skimmer dragonfly (*Plathemis lydia*) males have higher wing loading than females [105], which might relate to the speed required for male-male chases in this territorial species. Similarly, in wandering albatrosses, males have a higher wing loading than females and juveniles, which may make them better suited to the windier sub-Antarctic and Antarctic regions. The lower wing loading of adult females and fledglings may confer an advantage in exploiting the lighter winds of subtropical and tropical regions [106]. Of note, dimorphism is not universal as it has not been detected in vultures [107], fork-tailed flycatchers [108], or houseflies [109]. Our data does not allow us to determine whether differences in wing loading between sexes have a functional role.

Wing loading also varies within species. For example, *D. buzzati* highland populations show higher values of wing loading than lowland populations [110]. *Drosophila melanogaster* in the Indian continent shows the opposite pattern and shows a negative correlation between wing loading and latitude [111]. Wing loading and latitude are positively correlated in male and female warblers, suggesting intraspecific variation also occurs in vertebrates [112]. An expansion of our dataset that takes into consideration the effects of latitude on body size might reveal how wing and wing loading evolution affect biogeographical species distribution.

Microenvironmental conditions can also affect wing development and thus wing loading. Beetles fed on poor diets show substantially lower wing loading, which joint with poor general condition, ultimately precluded their ability to fly [101]. In pallid bats (*Antrozous pallidus*), wing loading depends on life stage [113]. Note, however, that this is not a consideration for insects, in which immature stages do not fly. No study has directly evaluated the effect of diet and crowding in *Drosophila,* but larval diet has effects on essentially all adult traits [114, 115]. While a robust body of literature has measured plasticity on mass and wing area, few studies have quantified the effect of environment on the integrated phenotype, wing loading. Given the importance of wing loading in species vagility, structure, and even for conservation potential, the time is ripe to understand the magnitude of plasticity in wing loading. A robust research avenue would address the effect of environmental conditions, such as crowding, diet, and temperature, on wing development and shape [for example, see 116].

Furthermore, time series can reveal the tempo and mode of evolution of wing morphology and wing loading. A museum specimen survey for the Iberian wasp (*Dolichovespula sylvestris*) revealed decreases in body size and wing area [117] but no change in wing loading over 100 years of sampling. Wing shape in the common Nightingale (*Luscinia megarhynchos*) has also changed, and become shorter, in the last 20 years in response to increased drought in the Iberian peninsula. These shorter wings are maladaptive for long migrations but might be beneficial for the survival of younglings in xeric conditions [118]. In a similar vein, experimental evolution can also reveal evolutionary patterns of wing morphology in organisms that can be reared in laboratory conditions. Experimental populations of *D. melanogaster* that undergo truncating selection on wing shape respond rapidly; nonetheless, the populations return to their original wing shape after selection is removed, potentially indicating strong constraints in wing shape from other correlated traits [119].

A second finding of our survey is the negative allometry in *Drosophila* wings. Wing area and wingspan in hoverflies show a negative allometric relationship with body mass [120]. Wing morphology in birds also shows a strong pattern of negative allometry in their wings [4]; however, this result is dependent upon the incorporation of phylogenetic information, as Rayner [2] showed positive allometry for both avian wing length and wing area using the same data that were subsequently reassessed by Taylor and Thomas [4]. This disparity underscores the importance of incorporating phylogenetic information in studies of allometric scaling. Our finding of negative wing size allometry is not universal [121]. Also in hoverflies, wing chord shows an isometric relationship indicating that there are differences in scaling factors among wing traits [120]. Similarly, wing scaling factors in bats are much closer to the isometric expectation than those of birds or insects. The wing area exponent is 0.69 for megabats and frugivorous microbats, whereas it is 0.60 for vespertilionids and 0.62 for molossids. This scaling differs between the three surveyed bat clades [121]. Comparative studies in insects have found that the isometric expectation is fulfilled at the interorder and intraspecific level but not at the level of intrafamily relationships in bees, where wing area increases hyperallometrically [122]. These differences indicate that interaction between wing area and mass differs across evolutionary dynamics of flight among groups that have evolved flight independently.

Our scoring scheme in a common environment allowed us to control for environmental variation by rearing all species under laboratory conditions, but still warrants consideration of potential limitations. The included species in our survey have vastly different range distributions. For example, *D. santomea* occurs only in the tropical island of São Tomé right on the Equator [123–125], and we also included *D. lummei*, which has a palearctic distribution [126, 127], and *D. melanogaster*, which is a cosmopolitan species that occurs around the world [128–130]. Future research should address the extent of trait divergence shown by wing loading and other traits that might affect the distribution and geographic range of *Drosophila* and other flying species.

Altogether, our results provide a roadmap for incorporating a phylogenetic perspective based on genome-wide variation to address classical questions in biomechanics. The ability to manipulate environmental conditions for *Drosophila* represents an open opportunity to understand how biomechanically-relevant traits can be studied from a genetic perspective.

## Author contributions

JAR and DRM conceived of the project

JAR and DRM designed the study

JAR, DRM, and GAJ collected data

JAR, PWK and DRM conducted analyses

JAR, and DRM wrote the initial manuscript draft

All authors edited the manuscript

## Acknowledgements

We thank the Matute lab for their helpful comments on this manuscript. Funding for this work was provided by the National Institute of Environmental Health Services (F32ES035271) to Jonathan A. Rader, the National Institute of Allergy and Infectious Disease (2T32AI052080) to Patrick W. Kelly, and the National Institute of General Medical Sciences (R35GM148244) to Daniel R. Matute.

## Data Accessibility Statement

The raw and processed egg images, along with our dataset of egg measurements and analysis scripts are archived at FIGSHARE (TBD upon acceptance).

