## Supplementary figures and table legends for "Body size, wing area, and wing loading follow a pattern of trait conservatism in *Drosophila*"

**SUPPLEMENT. RADER ET AL.**

#### SUPPLEMENTARY TABLES

Each table is presented in .csv format.

**TABLE S1. Species and isofemale lines used in this study.**

**TABLE S2. Comparative phylogenetics trait-evolution models used in this study.** BM: Brownian motion, OU: Ornstein–Uhlenbeck, EB: Early burst.

**TABLE S3.** Estimated species wing area means (emmean) by sex from ordinary least squares with standard errors (SE) and 95% confidence limits (lower.CL, upper.CL), and with degrees of freedom (df), t ratios (t.ratio) and p-values (p.value) from tests against 0.

**TABLE S4.**Estimated female-male differences (estimate) in mean wing area for each species from ordinary least squares with standard errors (SE) and 95% confidence limits (lower.CL, upper.CL), and with degrees of freedom (df), t ratios (t.ratio) and p-values (p.value) from tests against 0.

**TABLE S5.** Estimated pairwise interspecies differences (estimate) in mean wing area by sex from ordinary least squares with standard errors (SE) and 95% confidence limits (lower.CL, upper.CL), and with degrees of freedom (df), t ratios (t.ratio) and p-values (p.value) from tests against 0.

**TABLE S6.** Estimated species wing area means (emmean) by sex from phylogenetic generalized least squares with standard errors (SE) and 95% confidence limits (lower.CL, upper.CL), and with degrees of freedom (df), t ratios (t.ratio) and p-values (p.value) from tests against 0.

**TABLE S7.**Estimated female-male differences (estimate) in mean wing area for each species from phylogenetic generalized least squares with standard errors (SE) and 95% confidence limits (lower.CL, upper.CL), and with degrees of freedom (df), t ratios (t.ratio) and p-values (p.value) from tests against 0.

**TABLE S8.** Estimated pairwise interspecies differences (estimate) in mean wing area by sex from phylogenetic generalized least squares with standard errors (SE) and 95% confidence limits (lower.CL, upper.CL), and with degrees of freedom (df), t ratios (t.ratio) and p-values (p.value) from tests against 0.

**TABLE S9.** Estimated species wing area means (area.GM.fitted) by sex from Brownian Motion-based phylogenetic ANOVA (RRPP-BM; see Methods) with p-values (p.value) from tests against 0.

**TABLE S10.** Estimated pairwise interspecies differences (d) in mean wing area by sex, as well as estimated female-male differences (d) in mean wing area within each species, from Brownian Motion-based phylogenetic ANOVA (RRPP-BM; see Methods) with Upper Confidence Limits (UCL (95%)), Z scores, and P values (Pr > d) from tests against 0.

**TABLE S11.** Estimated species wing area means (area.GM.fitted) by sex from Kappa-based phylogenetic ANOVA (RRPP-Kappa; see Methods) with p-values (p.value) from tests against 0.

**TABLE S12.** Estimated pairwise interspecies differences (d) in mean wing area by sex, as well as estimated female-male differences (d) in mean wing area within each species, from Kappa-based phylogenetic ANOVA (RRPP-Kappa; see Methods) with Upper Confidence Limits (UCL (95%)), Z scores, and P values (Pr > d) from tests against 0.

**TABLE S13.** Estimated species body mass means (emmean) by sex from ordinary least squares with standard errors (SE) and 95% confidence limits (lower.CL, upper.CL), and with degrees of freedom (df), t ratios (t.ratio) and p-values (p.value) from tests against 0.

**TABLE S14.** Estimated female-male differences (estimate) in mean body mass for each species from ordinary least squares with standard errors (SE) and 95% confidence limits (lower.CL, upper.CL), and with degrees of freedom (df), t ratios (t.ratio) and p-values (p.value) from tests against 0.

**TABLE S15.** Estimated pairwise interspecies differences (estimate) in mean body mass by sex from ordinary least squares with standard errors (SE) and 95% confidence limits (lower.CL, upper.CL), and with degrees of freedom (df), t ratios (t.ratio) and p-values (p.value) from tests against 0.

**TABLE S16.** Estimated species body mass means (emmean) by sex from phylogenetic generalized least squares with standard errors (SE) and 95% confidence limits (lower.CL, upper.CL), and with degrees of freedom (df), t ratios (t.ratio) and p-values (p.value) from tests against 0.

**TABLE S17.** Estimated female-male differences (estimate) in mean body mass for each species from phylogenetic generalized least squares with standard errors (SE) and 95% confidence limits (lower.CL, upper.CL), and with degrees of freedom (df), t ratios (t.ratio) and p-values (p.value) from tests against 0.

**TABLE S18.** Estimated pairwise interspecies differences (estimate) in mean body mass by sex from phylogenetic generalized least squares with standard errors (SE) and 95% confidence limits (lower.CL, upper.CL), and with degrees of freedom (df), t ratios (t.ratio) and p-values (p.value) from tests against 0.

**TABLE S19.** Estimated species body mass means (mass.GM.fitted) by sex from Brownian Motion-based phylogenetic ANOVA (RRPP-BM; see Methods) with p-values (p.value) from tests against 0.
**TABLE S20.** Estimated pairwise interspecies differences (d) in mean body mass by sex, as well as estimated female-male differences (d) in mean body mass within each species, from Brownian Motion-based phylogenetic ANOVA (RRPP-BM; see Methods) with Upper Confidence Limits (UCL (95%)), Z scores, and P values (Pr > d) from tests against 0.

**TABLE S21.** Estimated species body mass means (mass.GM.fitted) by sex from Kappa-based phylogenetic ANOVA (RRPP-Kappa; see Methods) with p-values (p.value) from tests against 0.

**TABLE S22.** Estimated pairwise interspecies differences (d) in mean body mass by sex, as well as estimated female-male differences (d) in mean body mass within each species, from Kappa-based phylogenetic ANOVA (RRPP-Kappa; see Methods) with Upper Confidence Limits (UCL (95%)), Z scores, and P values (Pr > d) from tests against 0.

**TABLE S23.** Estimated species wing loading means (emmean) by sex from ordinary least squares with standard errors (SE) and 95% confidence limits (lower.CL, upper.CL), and with degrees of freedom (df), t ratios (t.ratio) and p-values (p.value) from tests against 0.

**TABLE S24.** Estimated female-male differences (estimate) in mean wing loading for each species from ordinary least squares with standard errors (SE) and 95% confidence limits (lower.CL, upper.CL), and with degrees of freedom (df), t ratios (t.ratio) and p-values (p.value) from tests against 0.

**TABLE S25.** Estimated pairwise interspecies differences (estimate) in mean wing loading by sex from ordinary least squares with standard errors (SE) and 95% confidence limits (lower.CL, upper.CL), and with degrees of freedom (df), t ratios (t.ratio) and p-values (p.value) from tests against 0.

**TABLE S26.** Estimated species wing loading means (emmean) by sex from phylogenetic generalized least squares with standard errors (SE) and 95% confidence limits (lower.CL, upper.CL), and with degrees of freedom (df), t ratios (t.ratio) and p-values (p.value) from tests against 0.

**TABLE S27.** Estimated female-male differences (estimate) in mean wing loading for each species from phylogenetic generalized least squares with standard errors (SE) and 95% confidence limits (lower.CL, upper.CL), and with degrees of freedom (df), t ratios (t.ratio) and p-values (p.value) from tests against 0.

**TABLE S28.** Estimated pairwise interspecies differences (estimate) in mean wing loading by sex from phylogenetic generalized least squares with standard errors (SE) and 95% confidence limits (lower.CL, upper.CL), and with degrees of freedom (df), t ratios (t.ratio) and p-values (p.value) from tests against 0.

**TABLE S29.** Estimated species wing loading means (load.GM.fitted) by sex from Brownian Motion-based phylogenetic ANOVA (RRPP-BM; see Methods) with p-values (p.value) from tests against 0.

**TABLE S30.** Estimated pairwise interspecies differences (d) in mean wing loading by sex, as well as estimated female-male differences (d) in mean wing loading within each species, from Brownian Motion-based phylogenetic ANOVA (RRPP-BM; see Methods) with Upper Confidence Limits (UCL (95%)), Z scores, and P values (Pr > d) from tests against 0.

**TABLE S31.** Estimated species wing loading means (load.GM.fitted) by sex from Kappa-based phylogenetic ANOVA (RRPP-Kappa; see Methods) with p-values (p.value) from tests against 0.

**TABLE S32.** Estimated pairwise interspecies differences (d) in mean wing loading by sex, as well as estimated female-male differences (d) in mean wing loading within each species, from Kappa-based phylogenetic ANOVA (RRPP-Kappa; see Methods) with Upper Confidence Limits (UCL (95%)), Z scores, and P values (Pr > d) from tests against 0.

**TABLE S33. wAIC comparisons to differentiate between three models (BM, OU, White noise) of trait evolution in wing area, body mass, and wing loading.** Table 2 shows wAIC values for an extended suite of evolutionary models of trait evolution.

**TABLE S34.**

### SUPPLEMENTARY FIGURES

**FIGURE S1.Heatmap with pairwise (x- versus y-axis) P-values (value and corresponding shading in each cell) for interspecies comparisons of wing area from an OLS model (see Methods).** NS = Not Significant at 0.05.

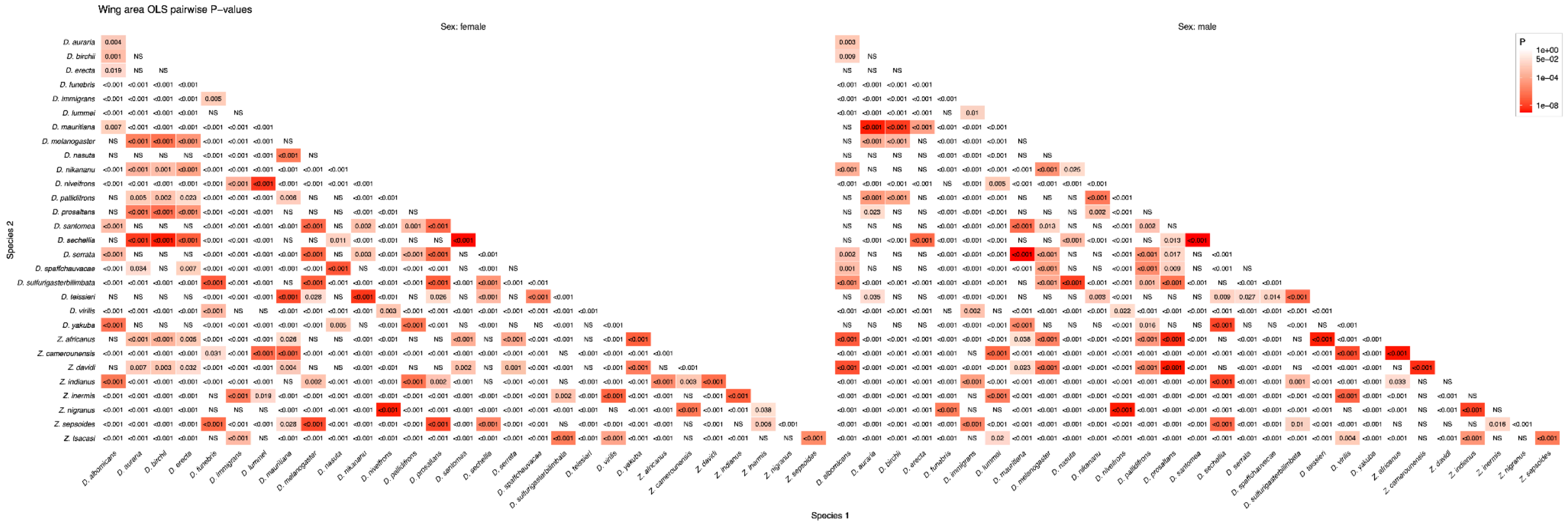

**FIGURE S2. Heatmap with pairwise (x- versus y-axis) P-values (value and corresponding shading in each cell) for interspecies comparisons of wing area from a PGLS model (see Methods).** NS = Not Significant at 0.05.

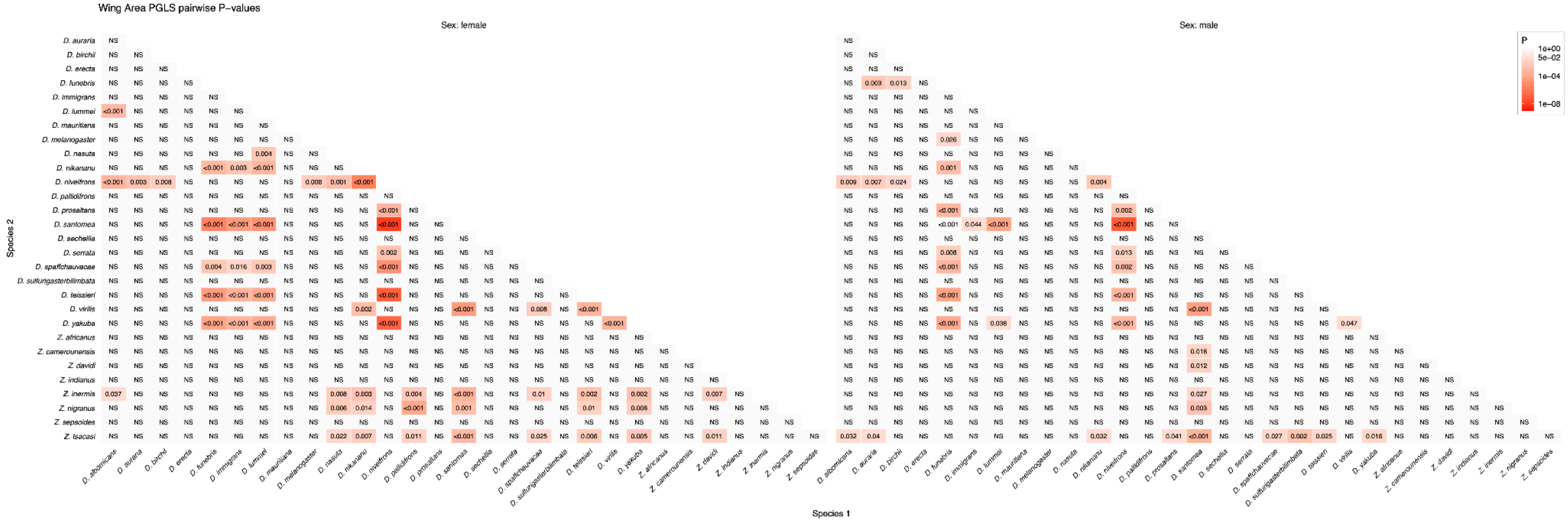

**FIGURE S3. Heatmap with pairwise (x- versus y-axis) P-values (value and corresponding shading in each cell) for interspecies comparisons of wing area from the RRPP-BM model (see Methods).** NS = Not Significant at 0.05.

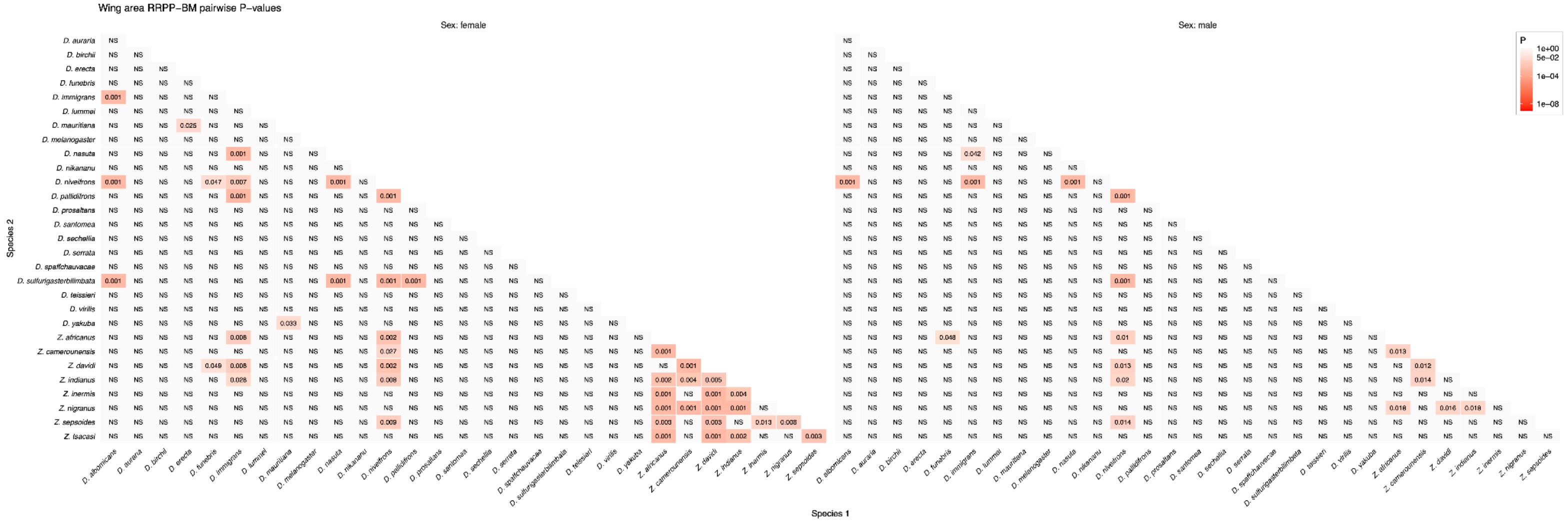

**FIGURE S4. Heatmap with pairwise (x- versus y-axis) P-values (value and corresponding shading in each cell) for interspecies comparisons of wing area from the RRPP_Kappa model (see Methods).** NS = Not Significant at 0.05.

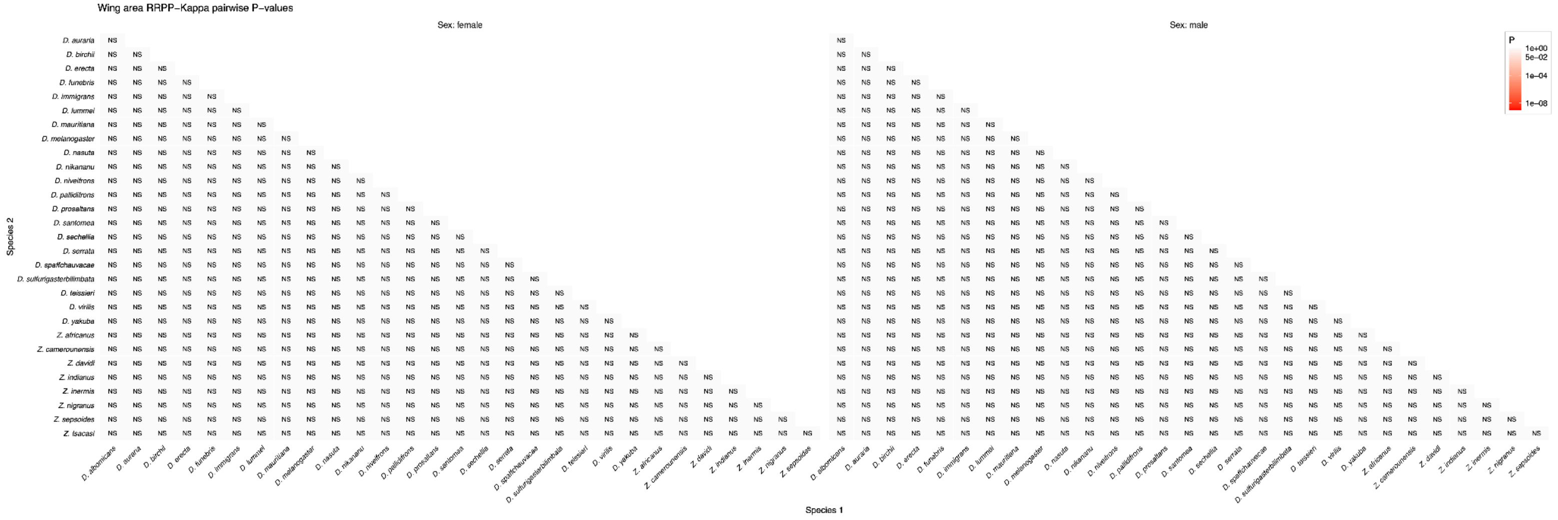

**FIGURE S5. Heatmap with pairwise (x- versus y-axis) P-values (value and corresponding shading in each cell) for interspecies comparisons of body mass from the OLS model (see Methods).** NS = Not Significant at 0.05.

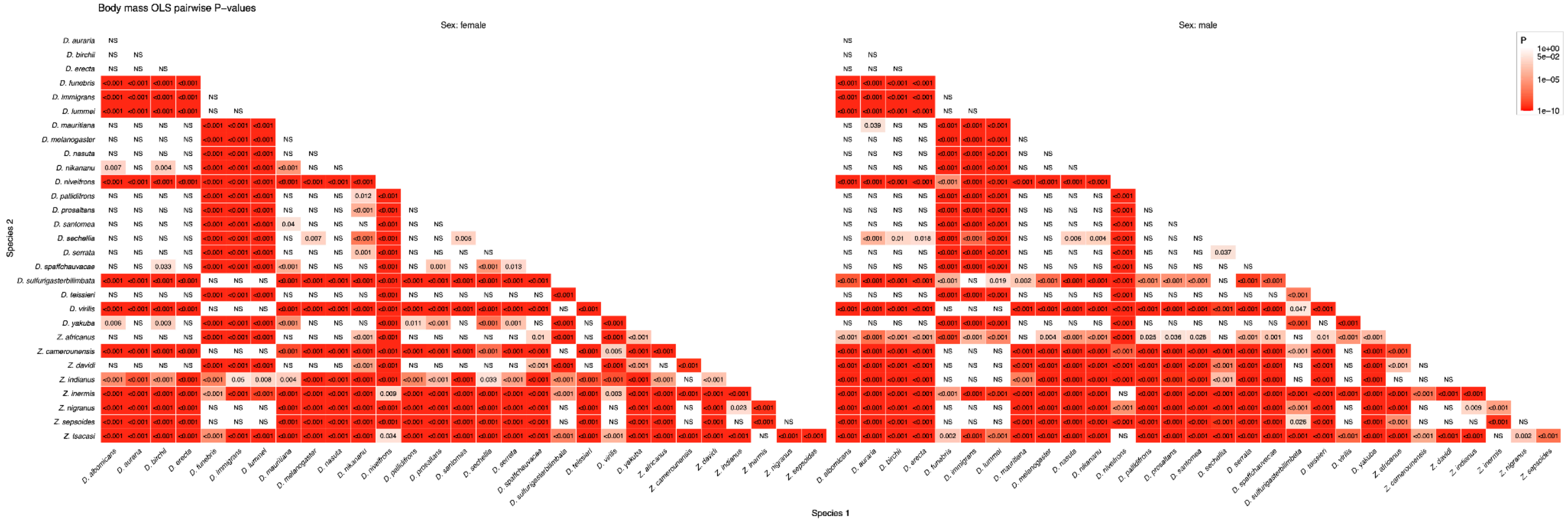

**FIGURE S6.Heatmap with pairwise (x- versus y-axis) P-values (value and corresponding shading in each cell) for interspecies comparisons of body mass from the PGLS model (see Methods).** NS = Not Significant at 0.05.

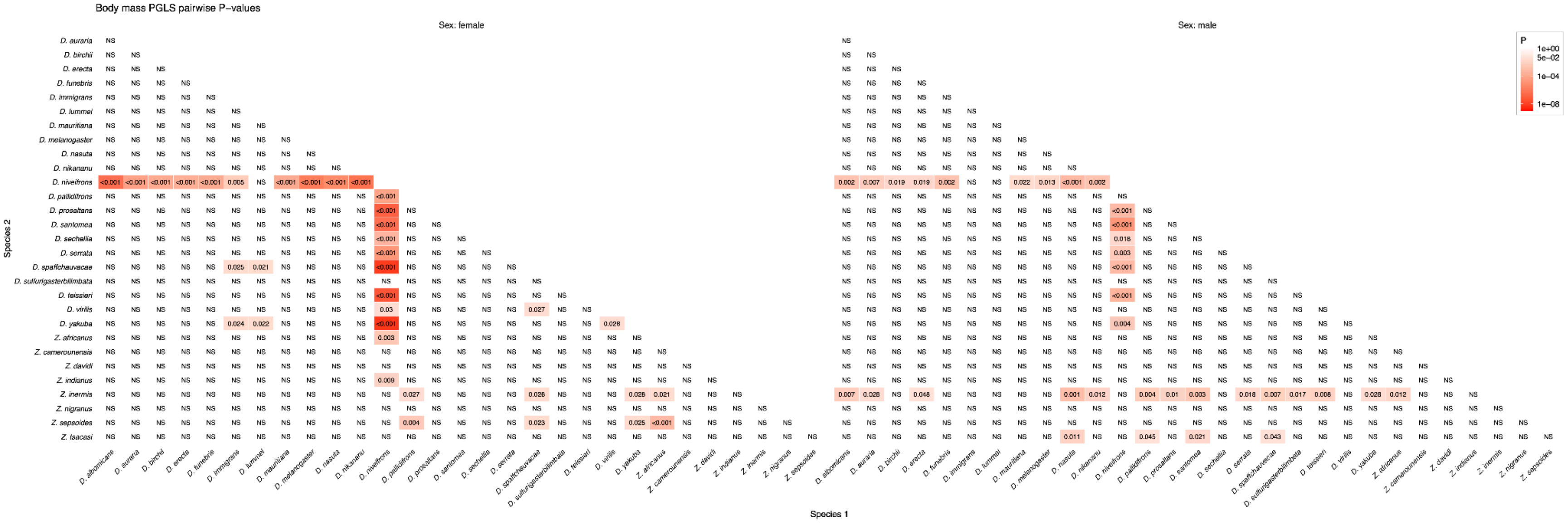

**FIGURE S7.Heatmap with pairwise (x- versus y-axis) P-values (value and corresponding shading in each cell) for interspecies comparisons of body mass from the RRPP-BM model (see Methods).** NS = Not Significant at 0.05.

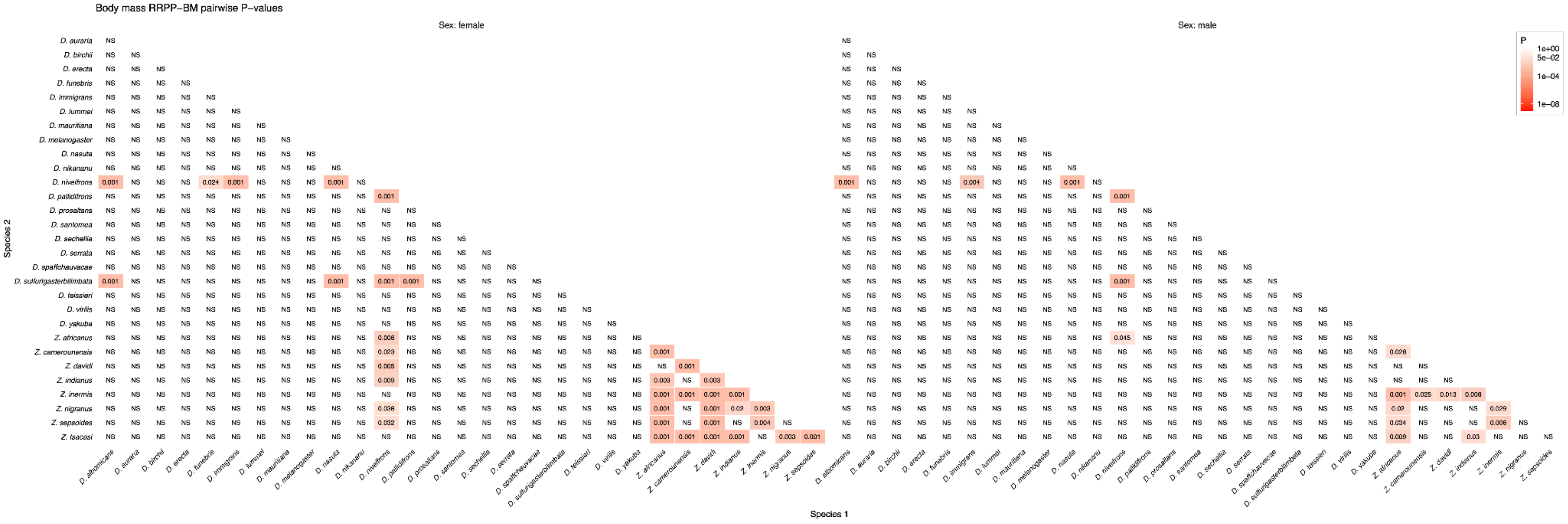

**FIGURE S8.Heatmap with pairwise (x- versus y-axis) P-values (value and corresponding shading in each cell) for interspecies comparisons of body mass from the RRPP-Kappa model (see Methods).** NS = Not Significant at 0.05.

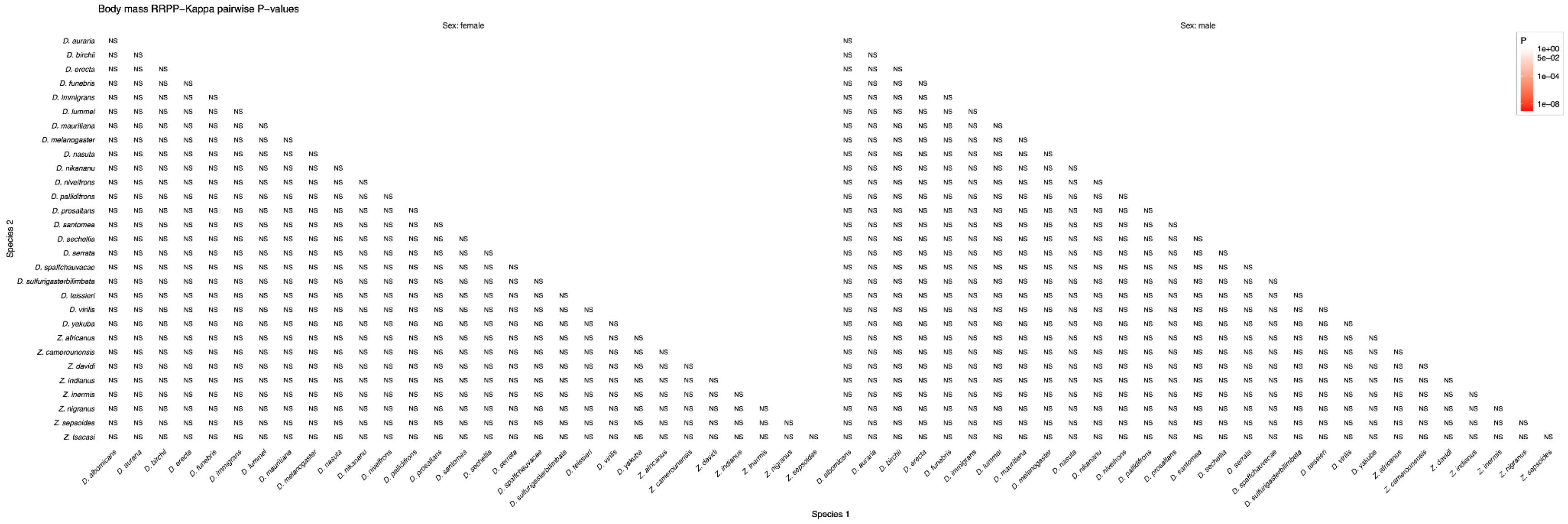

**FIGURE S9. Heatmap with pairwise (x- versus y-axis) P-values (value and corresponding shading in each cell) for interspecies comparisons of wing loading from the OLS model (see Methods).** NS = Not Significant at 0.05.

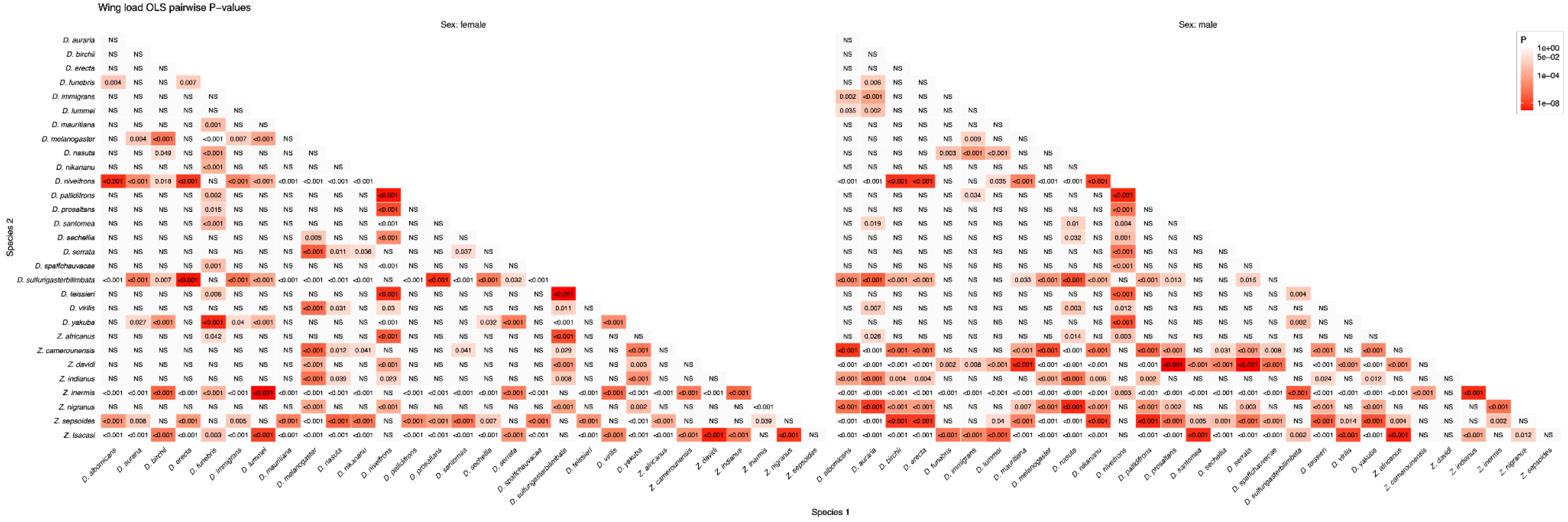

**FIGURE S10. Heatmap with pairwise (x- versus y-axis) P-values (value and corresponding shading in each cell) for interspecies comparisons of wing loading from the PGLS model (see Methods).** NS = Not Significant at 0.05.

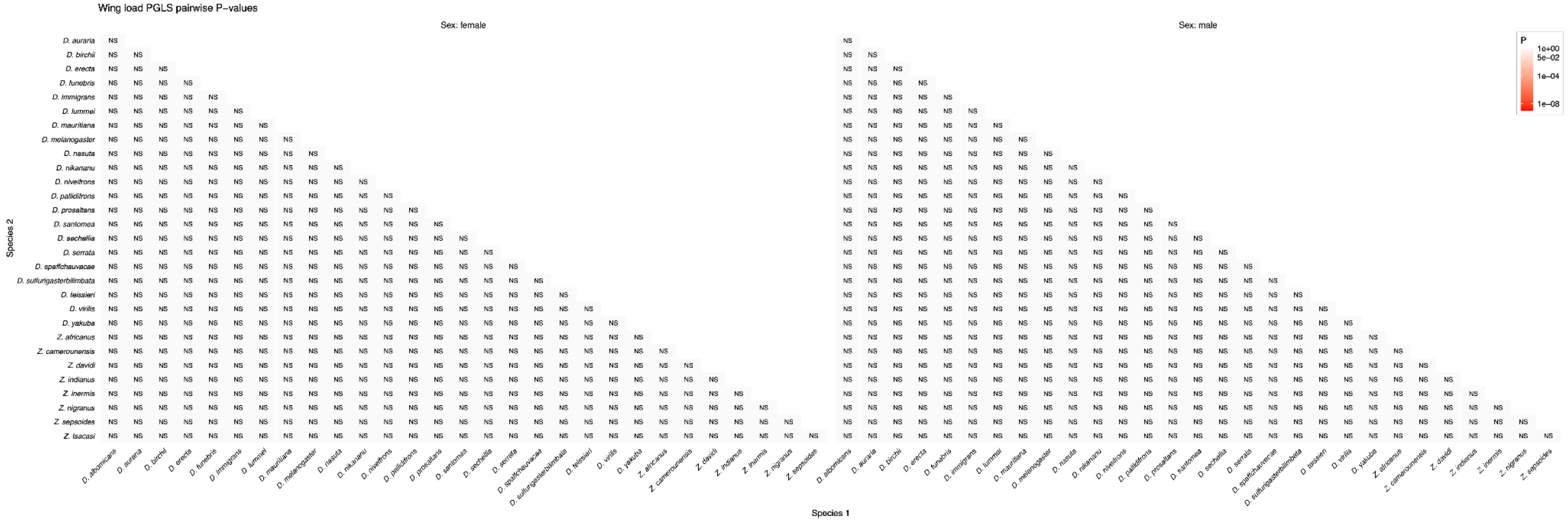

**FIGURE S11.Heatmap with pairwise (x- versus y-axis) P-values (value and corresponding shading in each cell) for interspecies comparisons of wing loading from the RRPP-BM model (see Methods).** NS = Not Significant at 0.05.

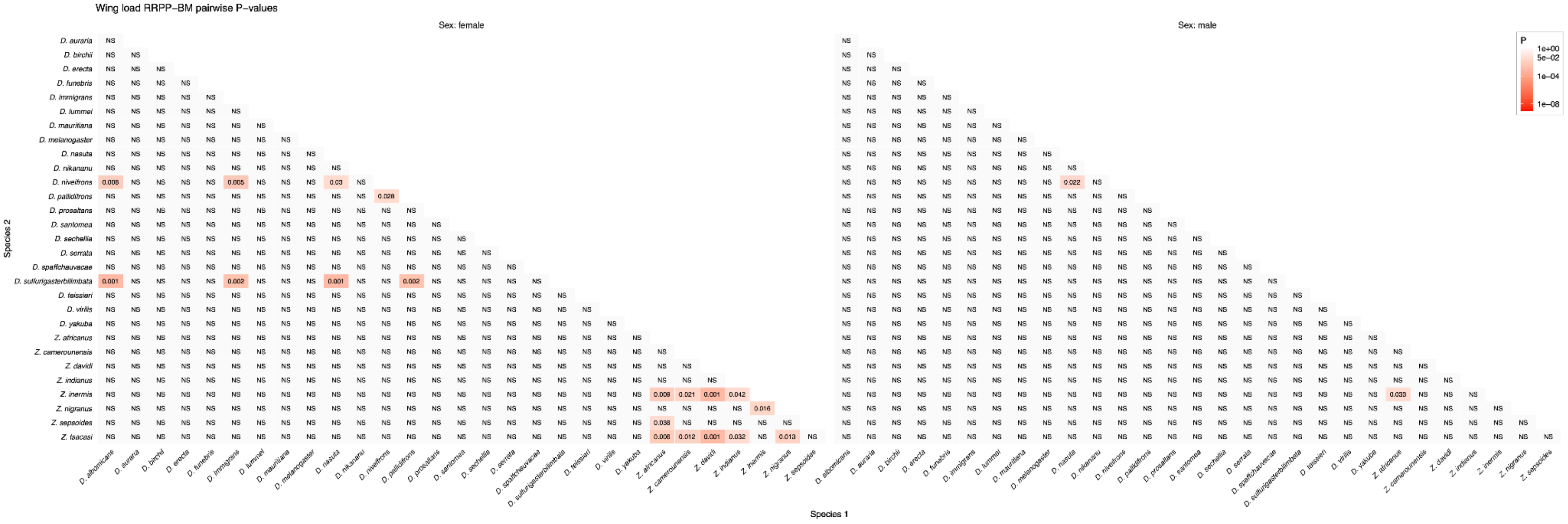

**FIGURE S12. Heatmap with pairwise (x- versus y-axis) P-values (value and corresponding shading in each cell) for interspecies comparisons of wing loading from the RRPP-Kappa model (see Methods).** NS = Not Significant at 0.05.

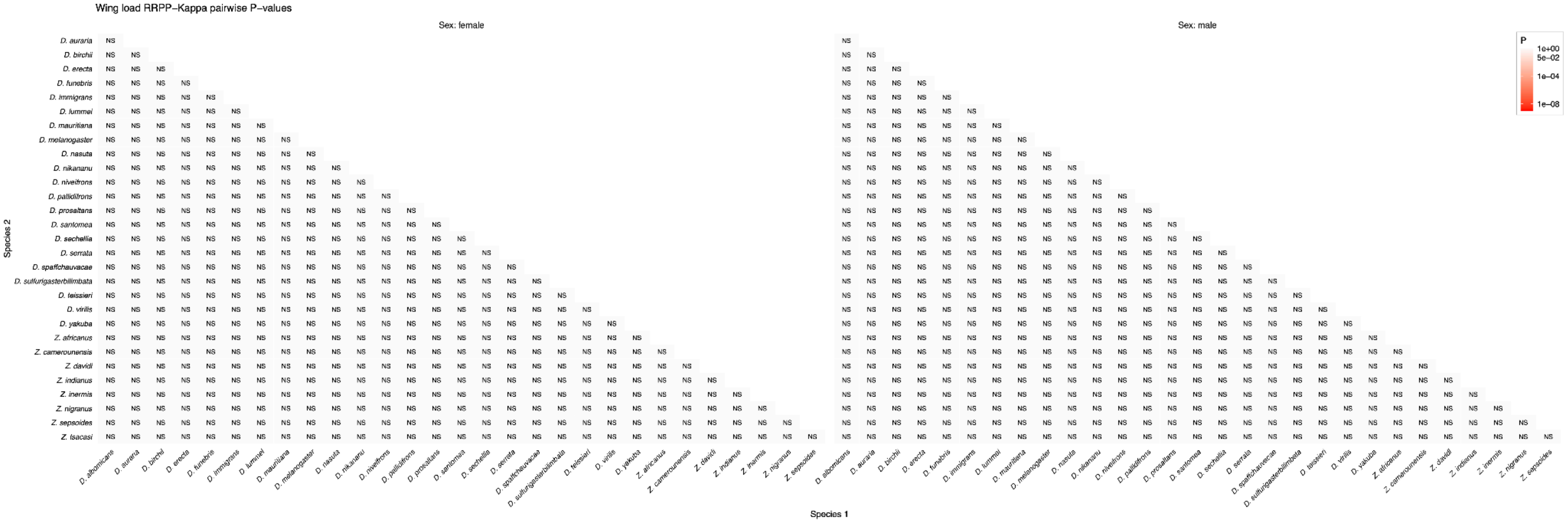
